# Intracellular phase separation of globular proteins facilitated by short cationic peptides

**DOI:** 10.1101/2021.07.08.450573

**Authors:** Vivian Yeong, Jou-wen Wang, Justin M. Horn, Allie C. Obermeyer

## Abstract

Phase separation provides intracellular organization and underlies a variety of cellular processes. These biomolecular condensates exhibit distinct physical and material properties. Current strategies for engineering condensate formation include using intrinsically disordered domains and altering protein surface charge by chemical supercharging or site-specific mutagenesis. We add to this toolbox by designing short, highly charged peptide tags that provide several key advantages for engineering protein phase separation. Herein, we report the use of short cationic peptide tags for sequestration of proteins of interest into bacterial condensates. Using a panel of GFP variants, we demonstrate how cationic tag and globular domain charge contribute to intracellular phase separation in *E. coli* and observe that the tag can affect condensate disassembly at a given net charge near the phase separation boundary. We showcase the broad applicability of these tags by appending them onto enzymes and demonstrating that the sequestered enzymes remain catalytically active.

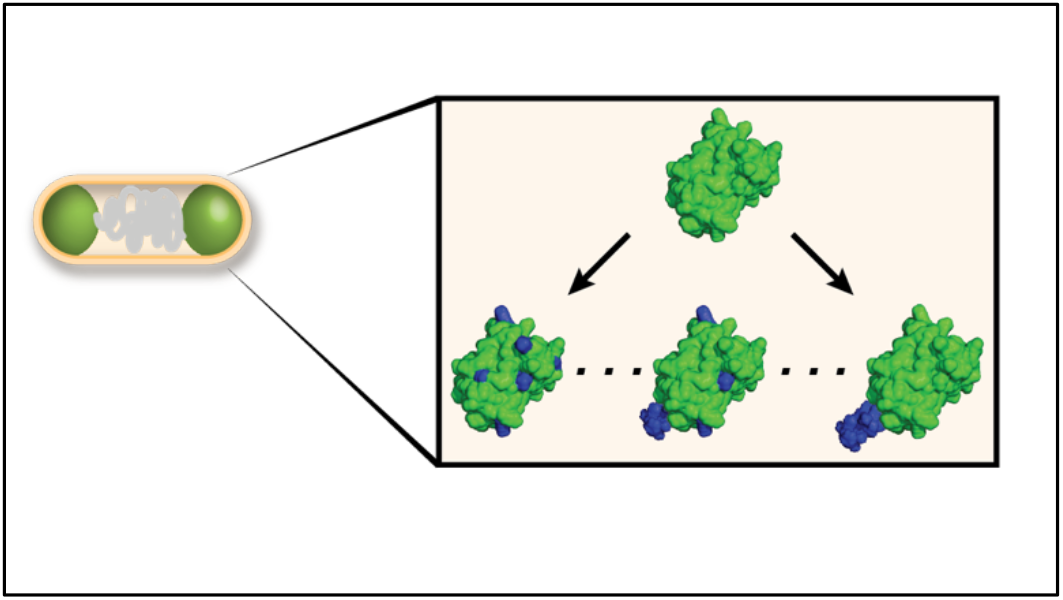

## MAIN

Biomacromolecules can undergo liquid-liquid phase separation to form membraneless entities, often referred to as biomolecular condensates^1^. These condensates organize cellular contents in a manner orthogonal to traditional membrane-delimited organelles. Biomolecular condensates have been shown to regulate diverse biological processes—for instance, cell signaling^2^, stress response^3^, and genome restructuring^4^—and exhibit many distinct properties. Condensates are dynamic, can maintain internal environments that are different from the surrounding cytoplasm, and have the ability to alter the folded structure and reaction kinetics of the macromolecules that are sequestered within them^1,5–9^. Consequently, there has been significant interest in imparting condensates with complementary responsive elements and enzymatic functionality. Recent developments include the ability to form and sequester proteins inside condensates in response to temperature, light, and small molecules, giving rise to engineered compartments with applications in synthetic biology and biocatalysis^6,10–16^.

Many native proteins that form biomolecular condensates contain intrinsically disordered regions (IDRs), which have been shown to undergo homotypic and heterotypic phase separation^17–20^. The fusion of these endogenous IDRs onto non-native globular proteins has enabled the formation of biorthogonal translation hubs, light-responsive metabolic clusters, and controlled release protein depots^13–15^. In addition to endogenous IDRs, engineered disordered sequences have also been used to promote intracellular phase separation^10,11^. For instance, the fusion of artificial IDPs to split proteins promotes recruitment to the condensed phase^11^. Moreover, phase behavior of these engineered IDPs can be tuned by altering the polypeptide sequence, primarily by controlling the relative aromatic and aliphatic content as well as molecular weight. While current strategies for engineering condensates take advantage of homotypic or specific biological, multivalent interactions to drive intracellular phase separation, the use of non-specific electrostatic interactions may provide a versatile alternative. Electrostatic interactions are a key driving force underlying the formation of many biomolecular condensates and have been shown to contribute to selective macromolecule partitioning in the condensed phase^5,17,19–21^. The IDRs of many proteins that natively phase separate are comprised of charged and aromatic residues^17,20^. Intermolecular interactions between these residues and other charged macromolecules in the cell are often sufficient to form a separate intracellular phase. Moreover, a number of studies have demonstrated that simply altering protein surface charge by genetic or chemical modification can also drive phase separation both *in vitro* and *in vivo*^22–25^.

Despite the success of current strategies to engineer synthetic condensates and protein sequestration, the tunability of these strategies remains a challenge and the ability to engineer multiple distinct condensates has not been established. Therefore, additional methods to promote intracellular phase separation are needed. While supercationic proteins have been demonstrated to form biomolecular condensates *in vivo*, global re-engineering of protein surface charge can be disruptive and difficult to execute. Very few native proteins are highly charged, and distributing charges on the protein surface requires a significant number of site-specific mutations (Fig. 1a)^26^. Consequently, the ability to supercharge a native protein by appending a small, highly charged tag offers several key advantages for promoting electrostatically-driven intracellular phase separation.

**Figure 1.**
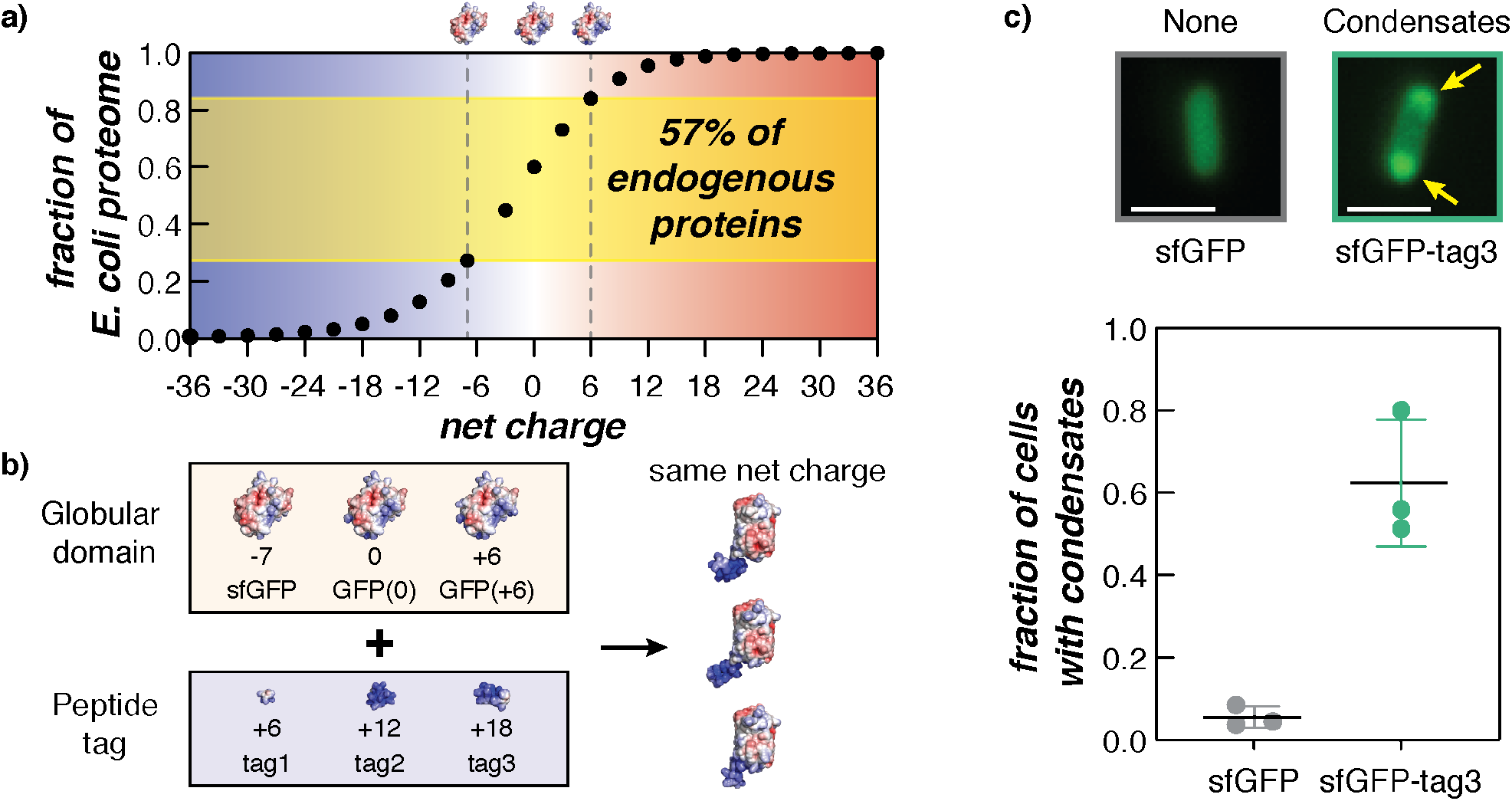
Cationic peptide tags promote the formation of biomolecular condensates in *E. coli*. (a) The cumulative distribution of overall net charge in the *E. coli* proteome reveals that the majority of proteins are not highly charged. 57% of proteins have an expected net charge between −7 and +6. (b) A schematic summarizing the GFP variants used in this study to understand how the protein globular domain and cationic peptide tags affect condensate formation in bacterial cells. (c) Microscopy images of *E. coli* cells expressing sfGFP (left) and sfGFP-tag3 (right) at 24 h post-induction. Intracellular condensates (indicated by ➙) are observed in cells expressing supercationic proteins. Scale bar is 2 μm. Images were analyzed using MicrobeJ and a custom MATLAB script to determine the fraction of cells that form condensates for each strain (n > 400 cells across three biological replicates per GFP variant). Each data point corresponds to a single biological replicate with at least 130 cells analyzed. The average and standard deviation of the three biological replicates are also plotted.

Although the use of small charge-dense tags may be advantageous for engineering protein phase separation, our understanding of how it alters the overall phase behavior of the resultant protein is limited. Several studies have investigated the role of charge anisotropy in the context of the charge patterning in polymeric systems and charge patchiness in native proteins; however, no studies have explored the effects of varying charge on both the phase separation domain and the protein globular domain^17,27–29^. A detailed examination of how both domains contribute to protein phase separation would provide additional insights for engineering biomolecular condensates in cells.

Here, we engineer intracellular condensate formation by appending to GFP short cationic peptide tags ranging from 9 to 27 amino acids in length. We investigate the interactions between these cationic peptides with various charged GFP globular domains that are representative of 57% of the *E. coli* proteome (Fig. 1a and 1b)^26^. We find that overall protein charge primarily governs intracellular phase separation; however, charge anisotropy may be important for tuning the persistence of condensates at a given charge threshold. We further establish the use of these cationic tags as a viable strategy for sequestering any protein of interest into bacterial condensates without the need to globally re-engineer protein surface charge or append large disordered phase separation domains.

## RESULTS

### Designing a library of GFPs with cationic peptide tags for intracellular phase separation

To investigate the contributions of the globular domain and peptide tag on bacterial condensate formation, we constructed a library of 9 GFP tagged variants that ranged in overall charge from 0 to +24 in increments of +6 (Fig. 1b and Supplementary Fig. 1). Each variant consisted of a charge-dense peptide tag appended to the C-terminus of an isotropically charged GFP—sfGFP(−7), GFP(0), or GFP(+6). Cationic peptide tags consisted of the amino acid sequence [GGSKKRKKR]_*n*_, whereby tag1 contained only GGSKKRKKR, tag2 contained two repeats of this sequence, etc. Since sfGFP had a net charge of −7, tagged sfGFP variants contained an additional lysine reside (+1) between the C-terminus of sfGFP and the [GGSKKRKKR]*n* tag in order to generate charge-equivalent variants (see the Supplementary Information). Previously reported untagged isotropic GFP variants^24,25^ were also included as controls to study the effects of overall positive charge and charge distribution on complex coacervation in *E. coli*.

Expression of GFP variants in *E. coli* was sufficient to induce condensate formation. Microscopy images acquired at 24 h post-induction depict the presence of GFP condensates in cells expressing sfGFP-tag3 and their absence in cells expressing the untagged control (Fig. 1c). Visual identification of condensates was verified by quantitative image analysis using MicrobeJ and a custom MATLAB script that calculated the fraction of condensate-containing cells detected in an image stack. Quantitative image analysis further confirmed that the majority of cells expressing the tagged GFP variant formed condensates.

### Parameters for condensate formation mediated by charge anisotropy

To examine the influence of the globular protein charge on cationic tag-driven coacervation, all GFP variants were expressed in *E. coli* and monitored for the formation of condensates by optical microscopy at 2, 4, 6, and 24 h post-induction (Supplementary Figs. 2-5). Broadly, cells expressing GFP variants with a net charge greater than +6 formed condensates at 24 h post-induction, suggesting that overall charge had a greater effect on intracellular phase separation than charge distribution (Fig. 2a). Moreover, this phase separation threshold is consistent with previous studies examining the intracellular phase separation of GFPs with isotropically-distributed surface charge^25^. Condensates were not observed in variants with a net charge ≤ +6 at 24 h. However, one notable exception was sfGFP-tag2, which formed weak condensates at 24 h despite having a net charge (+6) at this threshold. This result suggests that while overall protein charge is the dominant parameter governing intracellular phase separation, charge distribution may have a small but noticeable effect, particularly near the critical point for phase separation.

**Figure 2.**
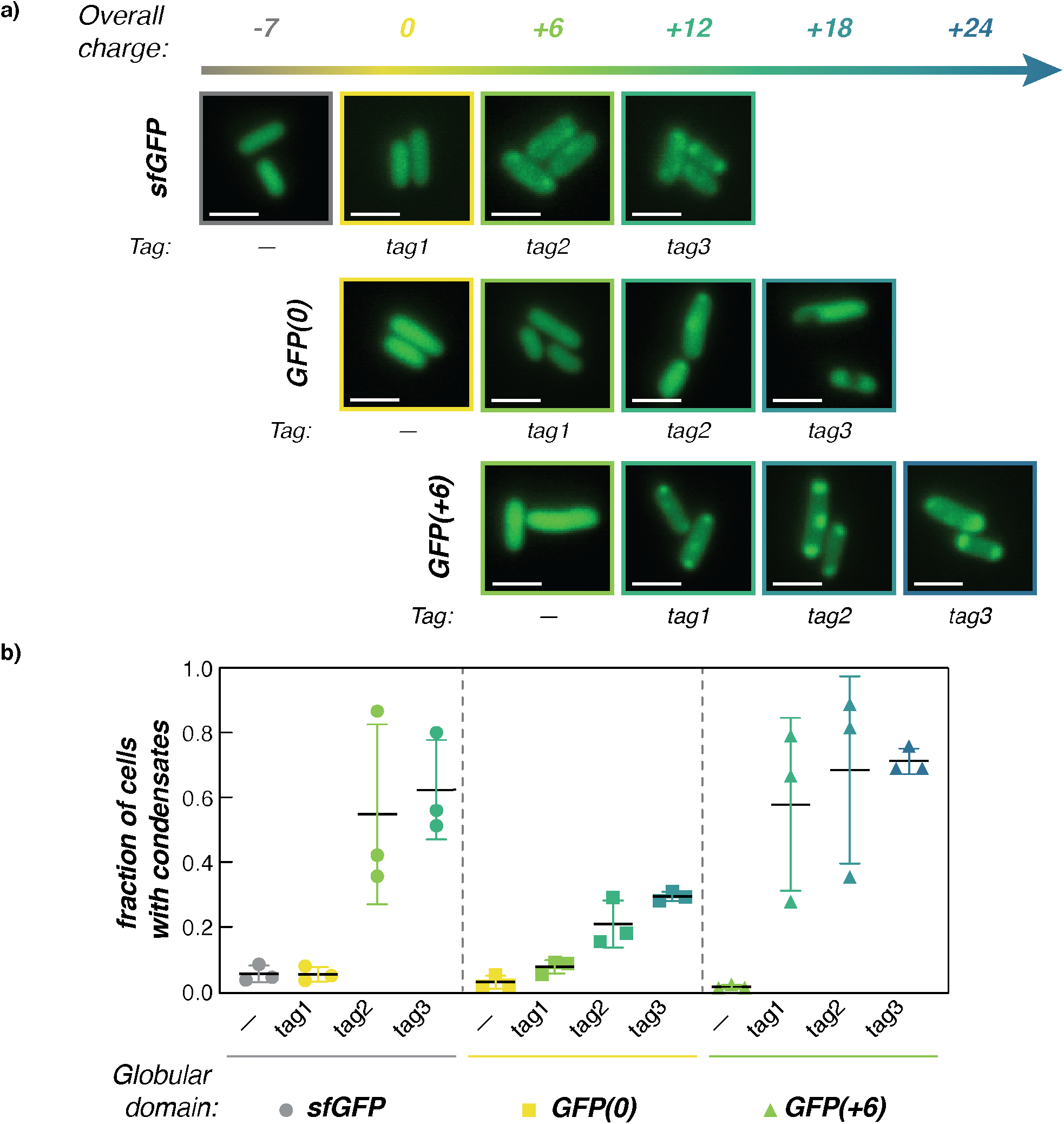
Intracellular phase separation depends on globular domain and peptide tag charge. (a) Representative microscopy images of *E. coli* cells expressing each GFP variant at 24 h post-induction. Generally, GFPvariants with an overall net charge ≥ + 12 formed intracelluar condensates. Additionally, sfGFP-tag2 (net charge +6) was also observed to form condensates at 24 h. Scale bar is 2 μm. (b) Analysis of n > 400 cells per GFP variant confirms that a net charge of ≥ + 12 resulted in condensate formation. Plot depicts the fraction of cells in which condensates were detected using an automated MATLAB script. The average and standard deviation of three biological replicates are plotted for each GFP variant.

Quantitative image analysis confirmed results from Fig. 2a and showed that condensates were not observed in GFP strains when the net charge was < +6 (Fig. 2b and Supplemental Figs. 6-7). While the automated image analysis did identify some false positives, the relatively low rate (< 0.11) enabled facile differentiation between strains that formed condensates and those that did not. To verify this approach, this method of analysis was also applied to previously reported isotropically-charged GFP variants as untagged controls, and revealed that cell fraction < 0.11 was an appropriate cutoff for determining the presence of condensates at 8 h and 24 h^25^ (Fig. 2b and Supplementary Fig. 6). Altogether, the results of this quantitative analysis fully aligned with those drawn from visual inspection.

### Traversing the phase boundary permits reversible condensate formation

A closer analysis of intracellular condensate formation over time revealed that while the majority of cationic GFP variants formed condensates that persisted, condensate disassembly was observed for two of the variants, both of which had an overall charge of +6. As a result, all GFP variants with a net charge of +6—untagged GFP(+6), GFP(0)-tag1, and sfGFP-tag2—were further examined and the fraction of cells containing condensates was plotted as a function of time (Fig. 3a). Similar plots for all untagged and tagged variants can be found in Supplementary Fig. 8.

**Figure 3.**
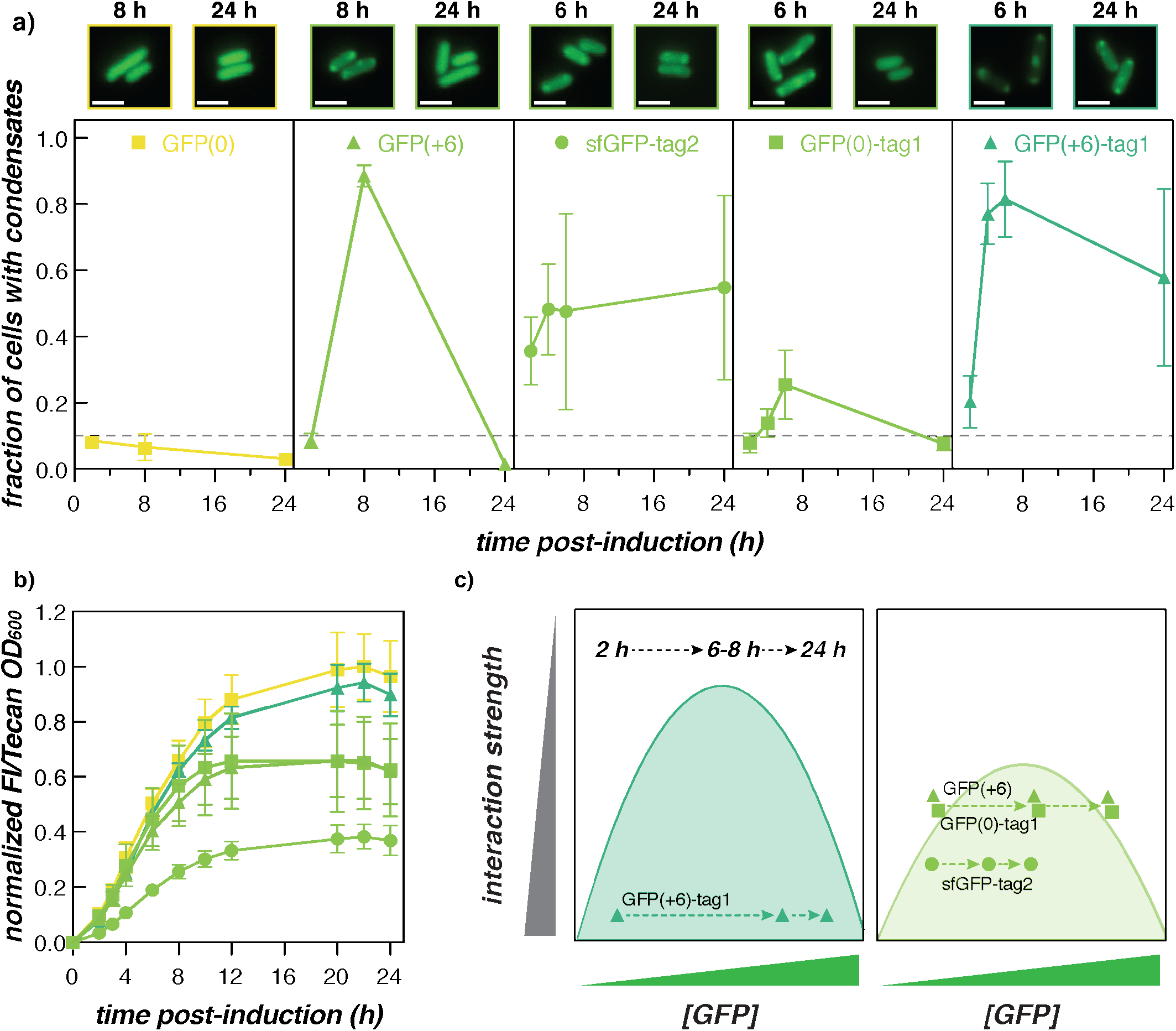
Changing GFP concentration allows for reversible intracellular phase separation. (a) The fraction of cells with condensates was monitored over time for select GFP variants. Interestingly, GFP(+6) and GFP(0)-tagl form condesates at 6-8 h, but not at earlier or later time points, suggesting that intracellular phase separation of supercationic GFP is reversible. Representative microscopy images of cells at the indicated time post-induction are provided (top). Scale bars are 2 μm. (b) The ratio of GFP fluorescence intensity to cell density was used as a proxy for intracellular GFP concentration. Intracellular concentrations of GFP(+6) and GFP(0)-tagl over time follow a similar trend that is distinct from the other GFP variants shown. The average and standard deviation of three biological replicates are plotted for each GFP variant. The legend is the same as for part a. (c) The schematic depicts how GFP(+6) and GFP(0)-tagl may undergo reversible condensate formation. The intracellular GFP concentrations and interaction strength of the cytoplasmic environment (i.e., the presence of other charged macromolecules and molecular crowders) lie along the phase boundary, enabling GFP(+6) and GFP(0)-tagl to traverse the phase boundary twice.

Reversible condensate formation has previously been observed in cells expressing GFP(+6) without a tag, whereby condensates formed at 8 h post-induction were completely disassembled by 24 h^25^. Here, we report that the charge-equivalent variant, GFP(0)-tag1, behaved similarly: condensates were first observed at 6 h and then fully disassembled at 24 h post-induction (Fig. 3a). This finding provides additional evidence that +6 may constitute a phase separation threshold for proteins at intracellular concentrations. As a control for globular domain charge, GFP(0) was not observed to phase separate at any time point, indicating that the inclusion of one tag repeat was responsible for the temporary formation of condensates in cells expressing GFP(0)-tag1. When appended to a more cationic globular domain, tag1 facilitated condensate formation at all time points (see GFP(+6)-tag1 in Fig. 3a). Interestingly, another charge-equivalent variant—sfGFP-tag2—formed condensates at 2 h that persisted until 24 h. Appending tag2 results in a larger increase in the charge patchiness on the protein, which may allow sfGFP-tag2 to undergo intracellular phase separation at both earlier and later time points.

We hypothesized that the reversible formation of condensates could be explained by increasing protein concentration throughout the phases of growth since phase separation is also a function of protein concentration^24,25,30^. Growth assays were performed for the 5 GFP variants highlighted in Fig. 3a, whereby optical density and GFP fluorescence intensity were monitored to approximate the intracellular GFP concentration over time (Fig. 3b and Supplementary Fig. 9). For all GFP variants, the ratio of GFP fluorescence to optical density increased exponentially from 2 to ~12 h post-induction and then plateaued. GFP(0) and GFP(+6)-tag1 demonstrated the highest GFP expression (Fig. 3b). In contrast, sfGFP-tag2 showed the lowest GFP expression with GFP(+6) and GFP(0)-tag1 exhibiting intermediate expression. In support of our hypothesis, GFP(+6) and GFP(0)-tag1 exhibited very similar GFP expression levels, suggesting that both charge-equivalent variants likely have similar underlying mechanisms for reversible phase separation. Since the 5 GFP variants exhibited different GFP concentrations at early (2 h), intermediate (6-8 h), and late (24 h) time points, we hypothesized that the mechanism for condensate reversibility involves traversing the intracellular binodal phase boundary and that this phase boundary varies with protein charge (Fig. 3c). We demonstrated that overall protein charge plays a more dominant role than charge distribution in determining intracellular phase separation (see Fig. 2). We speculate that this finding also applies broadly to the intracellular phase boundary and that changes in protein concentration appear to account for the observed differences in condensate behavior.

Previously, we demonstrated using isotropic GFP variants that the phase boundary broadens with increasing GFP charge when mixed with RNA *in vitro*^25^. We further showed that broadening of the phase boundary correlated with the propensity to form condensates in *E. coli*. In line with our previous findings, we hypothesize that GFP(+6)-tag1 exhibits a broader phase boundary and sufficient interaction strength, allowing the protein to remain phase separated despite a large change in GFP concentration during cell growth (Fig. 3c). In contrast, GFP(+6) and GFP(0)-tag1 have a lower net charge and, therefore, a smaller two-phase region in the cellular environment. This implies that the protein concentration in the cell traverses the phase boundary more than once to both form and disassemble condensates despite smaller changes in GFP concentrations. Finally, the charge-equivalent variant, sfGFP-tag2, is able to maintain condensate formation. The increased interaction strength provided by tag2 permits the protein to undergo intracellular phase separation at protein concentrations much lower than its charge-equivalent counterparts, and the small changes in protein concentration over time are not sufficient for traversal of the binodal phase boundary, allowing the condensate to persist. These experiments emphasize that sizeable perturbations can be made at the physiological phase boundary by modest modifications to protein charge anisotropy and concentration.

### Condensates behave as GFP/RNA coacervates

While some engineered bacterial condensates demonstrated reversible formation, the condensates formed by a majority of the GFP variants persisted. As result, engineered bacterial condensates were also investigated for physical properties characteristic of complex coacervates to establish that they were formed via these engineered electrostatic interactions. To probe the physical properties of the GFP condensates, solubility assays were conducted to demonstrate that condensates were held together by electrostatic interactions. GFP(+6)-tag3 was chosen as a representative tagged condensate-forming variant because it would be the least likely to undergo condensate reversibility in cells due to its high charge density. GFP(+36) was previously shown to be extracted from condensates using a buffer with high ionic strength and was used here as a positive control^25^. GFP(+6) was used as an untagged control that did not form persistent condensates. Protein expression was induced for 24 h and the presence or absence of condensates was verified by microscopy prior to cell harvesting. All cells were then lysed and washed using a buffer without NaCl (50 mM NaH_2_PO_4_, pH 8.0), and then resuspended in buffers with varying NaCl concentration (0, 0.1, or 1 M) (Fig. 4a). The fraction of GFP extracted from condensates was calculated by comparing the ratio of GFP relative to the total protein found in the soluble fraction as determined by SDS-PAGE analysis (Supplementary Fig. 10).

**Figure 4.**
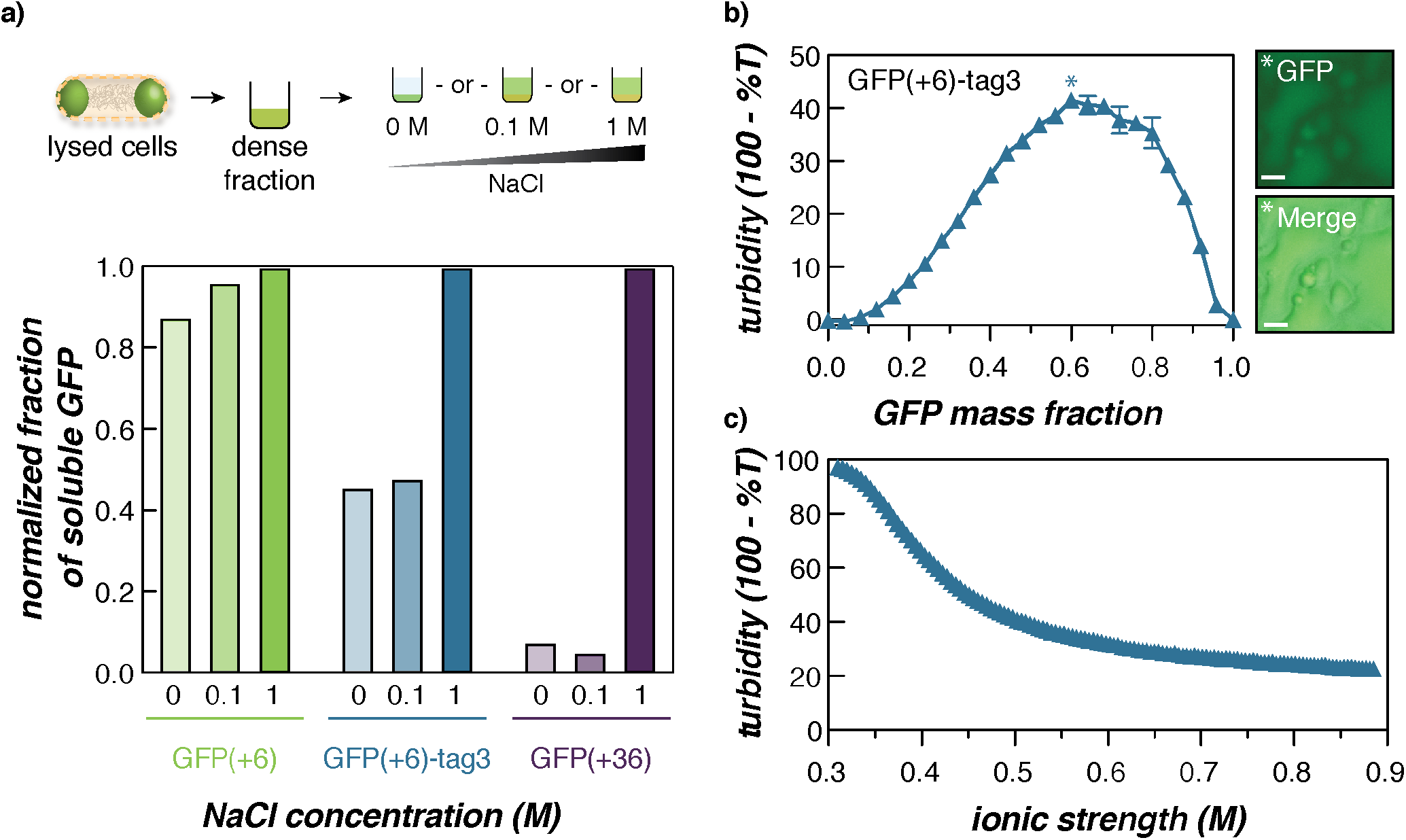
In vitro characterization of GFP(+6)-tag3/RNA coacervates. (a) Solubility of GFP(+6)-tag3 intracellular condensates in buffers of increasing ionic strength. Similar to the highly supercharged, condensate-forming variant—GFP(+36)— GFP(+6)-tag3 was extracted from condensates ex vivo by increasing NaCl concentration. (b) Turbidimetry measurements of GFP(+6)-tag3 with total RNA from torula yeast at 1 mg/mL total macromolecule concentration. Error bars represent the standard deviation of three technical replicates. Microscopy images (right) depict the GFP and merged (GFP with brightfield) channels of the GFP/RNA mixture at 0.6 GFP mass fraction, which demonstrated the highest turbidity. Scale bars are 10 μm. (c) NaCl titration was conducted at 0.6 GFP mass fraction in 1 mL total volume of GFP/RNA mixture prepared from 1 mg/mL protein and RNA stocks. Both turbidity and salt stability assays were conducted in a physiological buffer that mimics the intracellular ion concentration in E. coli (70 mM K_2_HPO_4_, 60 mM KCl, 40 mM NaCl, pH 7.4).

As expected, GFP(+6) showed no preference for extraction with different NaCl concentrations as it was soluble at the time cells were harvested (Fig. 4a). In contrast, the two condensate-forming variants— GFP(+6)-tag3 and GFP(+36)—preferred extraction in buffer supplemented with the highest NaCl concentration (1 M). Notably, a larger fraction of GFP(+6)-tag3 was extracted at lower NaCl concentrations when compared to GFP(+36). This can be explained by differences in overall net charge; since GFP(+6)-tag3 has a lower net charge (+24), its extraction requires less salt and, therefore, becomes more soluble at lower NaCl concentrations. Nonetheless, increased extraction of GFP(+6)-tag3 from condensates at higher ionic strength suggests the key role of electrostatics in the formation of condensates, which is consistent with the formation of complex coacervates.

To provide further evidence that condensates are formed through electrostatic interactions, we investigated the ability of GFP(+6)-tag3 to undergo phase separation *in vitro* with RNA, the most abundant anionic biopolymer in the cell by weight^31^. Purified GFP(+6)-tag3 was mixed with total RNA from torula yeast at a constant total macromolecule concentration in a buffer (70 mM K_2_HPO_4_, 60 mM KCl, 40 mM NaCl, pH 7.4) that mimicked intracellular ion concentrations and pH (Fig. 4b). Phase separation was observed across a wide range of mixing ratios under physiological conditions and a modest total macromolecule concentration. Moreover, the GFP/RNA mixture formed liquid-like coacervates that coalesced even at the mixing ratio corresponding to the highest turbidity (Fig. 4b and Supplementary Fig. 11). A NaCl titration was also performed to determine the critical salt concentration, which aligned with the results of the solubility assay, with the coacervate phase dissolving at an ionic strength of ~0.4 M (Fig. 4c). We note the discrepancy in the maximum turbidity value between the turbidity (Fig. 4b) and salt stability (Fig. 4c) assays was due to differences in the pathlength of the vessels in which measurements were obtained. Taken together, these *in vitro* experiments provide evidence for the complex coacervate-like nature of the engineered condensates formed from cationically tagged proteins in bacteria.

### Sequestration of enzymes in engineered bacterial condensates

We then wanted to demonstrate the broad utility of these cationic tags by examining how they affected the phase separation of a diverse set of proteins. Tags with two or three repeats were appended to mScarletI, a red fluorescent protein that has an expected charge of −3. Figure 5a depicts representative microscopy images of cells expressing the untagged and tagged mScarletI variants at 24 h post-induction. Addition of either tag was sufficient to drive condensate formation in cells. Quantitative image analysis revealed that while both tagged mScarletI variants remain phase separated at all time points, condensates become significantly less distinct for mScarletI-tag2 at 24 h (Supplementary Figs. 12-15).

**Figure 5.**
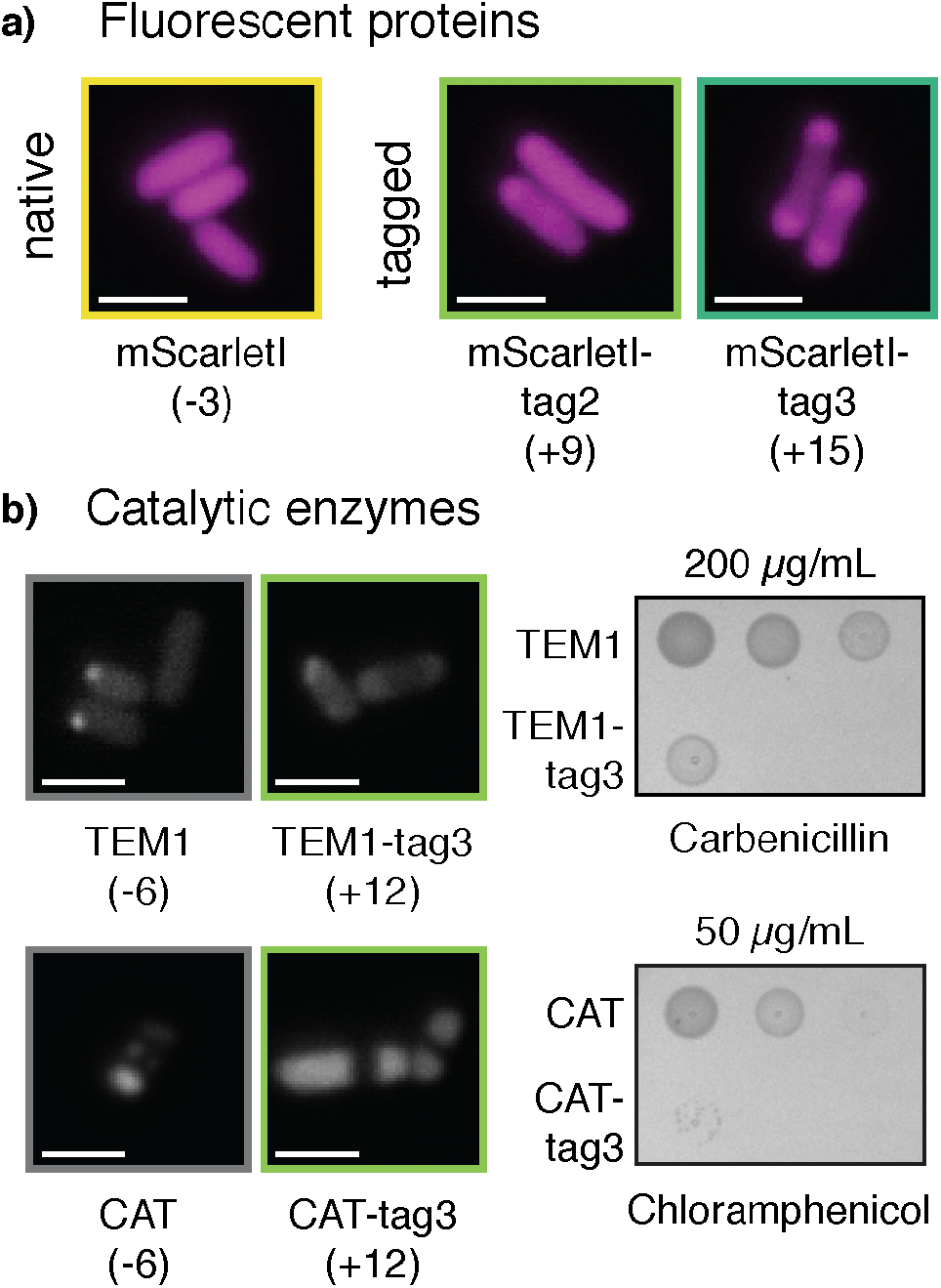
Cationic tags can induce the intracellular phase separation of fluorescent proteins and enzymes without abolishing catalytic activity. (a) Microscopy images of mScarletI and tagged variants at 24 h post-induction. Expected net charges are denoted in the parenthesis below. Appending cationic tag2 or tag3 promotes intracellular phase separation. (b) Microscopy images of beta lactamase (TEMI), chloramphenicol acetyltransferase (CAT), and each of their tagged derivatives at 24 h post-induction. Each protein also contained a N-terminal TC tag for fluorescence imaging with FlAsH/ReAsH (left). Scale bars on all micrographs are 2 μm. Spotting assay on LB plates supplemented with the respective antibiotic indicate that the presence of the cationic tag does not completely inhibit growth (right). Reduced fitness of the tagged relative to the untagged strains was also observed on control plates with no antibiotic (Supplementary Fig. 19).

To further validate the ability of cationic tags to promote the formation of biomolecular condensates, cationic tags were appended to enzymes that confer antibiotic resistance, TEM (β-lactamase) and CAT (chloramphenicol acetyltransferase) which degrade carbenicillin and chloramphenicol, respectively. Both native enzymes have an expected charge of −6, which is in line with the charge of the globular domains investigated using GFP variants. In addition to the cationic tag, a N-terminal tetracysteine (TC) tag was added to these proteins to enable visualization via fluorescence microscopy with FlAsH-EDT_2_ or ReAsH-EDT_2_. Staining unexpectedly revealed local regions of high protein concentration in cells expressing untagged TEM1 and CAT at 24 h post-induction; however, appending tag3 resulted in the formation of condensates that were similar in size to those formed with tagged fluorescent proteins (Fig. 5b and Supplementary Fig. 16). Moreover, the condensates observed were similar in size and morphology to sfGFP-tag3 condensates that were stained with ReAsH-EDT_2_, indicating that CAT-tag3 and TEM1-tag3 condensates were not an artifact of the staining procedure used (Supplementary Fig. 17).

Growth assays were also conducted to test the ability of untagged and tagged strains to survive on LB plates supplemented with the respective antibiotic and inducer (Supplementary Fig. 18). Both TEM1-tag3 and CAT-tag3 were able to grow on respective antibiotic plates (Fig. 5b). However, a clear reduction in fitness was observed in the absence of antibiotic (this was also observed for a sfGFP-tag3 control; Supplementary Fig. 19). In addition to arabinose, TEM1 variants were also grown on antibiotic plates supplemented with glucose, which reduces expression below uninduced levels. When grown on glucose plates without antibiotic, minimal growth differences were observed between the TEM1 variants, suggesting that condensate formation is responsible for the reduction in fitness. In spite of this, TEM1-tag3 maintained growth across all carbenicillin concentrations tested. CAT-tag3 also grew on 25 μg/mL and 50 μg/mL chloramphenicol plates but appeared to be more sensitive to higher concentrations of chloramphenicol. However, these differences in antibiotic resistance for CAT-tag3 and TEM1-tag3 variants cannot be directly compared due to variations in experimental setup (different inducible constructs were used to express the CAT and TEM1 variants resulting in varying levels of protein expression) and recommended antibiotic concentrations^32^ (see Supplementary Fig. 18). Although a reduction in fitness was observed for all tagged variants, incorporation of the cationic peptide tag allowed for increased sequestration of the enzyme into bacterial condensates without abolishing enzymatic activity.

## DISCUSSION

Herein, we report the ability of short cationic peptides to promote the formation of complex coacervate-based biomolecular condensates in *E. coli*. In this study, cationic tags of varying lengths were appended onto globular domains in order to investigate how the net charge of these individual components act in concert to affect intracellular phase separation. The expected charge on globular domains tested ranged from −7 to +6, which spans the expected charge of 57% of the *E. coli* proteome^26^. The use of 9-27 amino acid tags described here provides a simple and versatile approach to engineer protein phase separation in cells. We show that the shortest tag is able to promote condensate reversibility at the physiological phase boundary and that near the physiological phase boundary, tag length can determine condensate behavior.

Our results indicated that overall charge was the main parameter governing condensate formation as GFP variants with net charge > +6 generally phase separated regardless of the identity of the globular domain or tag. Even at high intracellular protein concentrations, GFP variants did not form condensates if they were not sufficiently charged. This is illustrated by the fact that GFP(+6)-tag1 formed condensates while GFP(0) did not even though both proteins exhibited similar intracellular protein concentrations. Moreover, we investigated the physical properties of condensates and verified *in vitro* that, like complex coacervates, they are held together by electrostatic interactions and are sensitive to the ionic strength of the solution.

While overall charge had a larger effect on condensate formation than charge distribution, the impact of charge distribution on phase separation became markedly pronounced near the phase separation threshold. Closer examination of GFP variants with a net charge of +6 revealed that the variants with increased charge patchiness (in particular, sfGFP-tag2) maintained condensates over time while the lower charge density variants formed condensates that disassembled at later time points. Various studies in protein and polypeptide systems have reported that increasing the charge anisotropy provides broader phase boundaries^27,29,33,34^, which we hypothesize allows sfGFP-tag2 to remain phase separated even as protein concentration and cytoplasmic interaction strength fluctuates. In contrast, lower charge density variants with a net charge of +6 have an anticipated decreased interaction strength and are closer to the critical point. As a result, these variants are able to traverse this phase boundary in response to changes in the cytoplasmic environment.

Cationic peptide tags are a useful tool for bioengineering and biological study given the dynamic nature of condensates, the ability to design condensate reversibility or persistence, and the ease of altering a protein’s overall charge density. The primary sequence of a globular protein can be used to predict the tag length/charge required for intracellular phase separation and appending these designer tags onto enzymes can result in their sequestration in condensates. Moreover, engineered enzymes within the coacervate-like condensate retain their activity, highlighting the broad applicability of these tags. Simultaneously, however, this approach may present a few limitations that require further characterization and improvement. Most critically, additional studies are needed to probe the impact of condensate formation on cell viability and methods to minimize growth defects are necessary in order to fully harness the utility of these cationic tags. The effects of these tags on very large proteins and recombinant proteins that have a tendency to aggregate are unknown. Additionally, we are limited by the protein expression system to a certain intracellular protein concentration regime. Enhanced control of protein expression would permit more precise delineation of the intracellular phase boundary. In addition, exploring the charge patterning of peptide tags may allow for further fine-tuning of condensate formation at or near the intracellular phase boundary. Striking an intricate balance between overall protein charge and charge anisotropy may allow for improved control of engineered condensate formation and disassembly. With this initial demonstration, we have expanded the groundwork for rational, charged-based engineering of condensate formation and protein sequestration. As an extension of this work, we envision using cationic peptide tags to recruit multiple proteins to intracellular condensates and build complex, functional organelles by incorporating enzymes involved in biochemical pathways.

## METHODS

### Cloning

Three different cationic tags (tag1, tag2, and tag3) were created by modifying the number of repeats of the amino acid sequence, GGSKKRKKR. These tags were then appended to each of the isotropic GFP (sfGFP(−7), GFP(0), and GFP(+6)) variants using a combination of PCR, restriction enzyme digestion, and T4 ligation. Primer sequences, plasmid templates, and restriction sites used can be found in the Supplementary Information along with details for cloning of mScarletI, CAT, and TEM1 variants. All GFP and mScarletI variants contained a N-terminal 6xHis tag unless specified otherwise.

### Protein Expression

Glycerol stocks of NiCo21(DE3) cells (New England Biolabs) transformed with each GFP variant were streaked onto LB agar plates with added ampicillin (100 μg/ml, Gold Biotechnology). A single colony was inoculated into sterilized LB media supplemented with ampicillin (100 μg/mL) and grown in an incubator (Thermo Scientific MaxQ 6000) overnight at 37 °C with shaking at 225 rpm. The overnight cultures were back diluted to OD_600_ ~ 0.1 in 25 mL LB media supplemented with ampicillin (100 μg/mL) in sterile 125 mL Erlenmeyer flasks. Cultures were grown at 37 °C, shaking at 225 rpm for 2-3 h. At OD_600_ = 0.8 −1.0, cultures were induced with 1 mM isopropyl β-D-1-thiogalactopyranoside (IPTG, Gold Biotechnology) and then kept at 25 °C with shaking at 225 rpm for 24 h. 20 μL aliquots were taken from the cultures at 0, 2, 4, 6, and 24 h after induction for imaging via optical microscopy.

### Optical Microscopy

Cell samples were applied to agarose pads for imaging. The agarose pads were made by preparing 1 w/v% agarose (TopVision) in milliQ water. 50 μL of melted agarose was then pipetted onto a 25 mm x 75 mm microscope slide and immediately covered by a 18 mm x 18 mm coverslip (ThermoScientific). After the agarose solidified, the coverslip was slowly removed and then 1.5 μL of cell culture was added on top of the agarose pad. The agarose pad with sample was gently covered with the coverslip and sealed on all four sides with clear nail polish. Cells were imaged on agarose pads at 0, 2, 4, 6, and 24 h post-induction. Images were taken using a 100X oil 1.40 NA UPlanSAPO objective (Olympus) with illumination by GFP (*λ*_*ex*_ = 479-522 nm; *λ*_*em*_ = 525-550 nm; EVOS GFP light cube) or Texas Red (*λ*_*ex*_ = 585-629 nm; *λ*_*em*_ = 628-632 nm; EVOS Texas Red light cube) and brightfield channels on an EVOS FL Auto 2 inverted fluorescent microscope. 10-16 images were taken for each sample in order to acquire a sufficient number of cells per strain (> 400 across three biological replicates).

### Image analysis

The microscopy images were pre-processed using ImageJ with a plug-in, MicrobeJ (version 15.3l (14)). Briefly, images were compiled into stacks and were subjected to a rolling ball background subtraction with a 100 pixel radius. Stacks of images were individually thresholded using the Li method in MicrobeJ and cells that met specific area, length, and width, circularity, and angularity constraints were identified and considered for analysis. Images of individual cells identified by MicrobeJ were loaded into MATLAB for processing and identification of condensates. A custom MATLAB script was used to identify condensates and determine the fraction of condensate-containing cells. The fraction of cells containing condensates was then plotted in GraphPad Prism (version 9.1.0). The specific constraints used and complete MATLAB code can be found in the Supplementary Information.

### Chloramphenicol Sensitivity Spotting Assay

Overnight cultures of TC-CAT, TC-CAT-tag3, TC-sfGFP, and TC-sfGFP-tag3 were prepared as described above and then diluted to OD_600_ = 0.1 in LB media supplemented with 100 μg/mL ampicillin and 10 μM FlAsH-EDT_2_ (Fisher Scientific). Cultures were grown at 37 °C with shaking at 225 rpm until they reached OD_600_ = 0.8 – 1, at which point cultures were induced with 1 mM IPTG and allowed to grow at 25 °C with shaking for 2 h. Subsequently, all cultures optical densities were normalized to OD_600_ = 1 in LB media supplemented with 100 μg/mL ampicillin. Normalized density cultures were diluted 100-fold into LB media. From this dilution, cultures were serially diluted 20-fold in LB media. 5 μL of all normalized cultures and subsequent dilutions (100X and 2,000X) were then spotted onto LB agar plates supplemented with 1 mM IPTG, 100 μg/mL ampicillin, and varying concentrations of chloramphenicol (0, 25, 50, 200, or 300 μg/mL). Samples on plates were allowed to dry fully before incubation in the dark at 25 °C for ~30 h. Plates were then imaged on a Gel Doc XR+ System (Bio-Rad). Colonies from the plate were also picked and diluted in 20 μL LB media and imaged on a microscope slide to confirm the presence of condensates (as described in “Optical Microscopy”). Spotting assays were similarly performed for TEM1 variants and additional details about these assays can be found in the Supplementary Information.

## DATA AND CODE AVAILABLILITY

The MATLAB code used to identify condensate-containing cells can be found in the Supplementary Information. All data is available upon request.

## ACKNOWLEDGEMENTS

We acknowledge the National Science Foundation (DMR: 1848388) for funding.

## AUTHOR CONTRIBUTIONS

V.Y., J.W., and A.C.O. conceived of the project. V.Y. and J.W. performed *in vitro* and *in vivo* experiments. V.Y., J.M.H., and A.C.O analyzed the data. V.Y. and A.C.O. wrote the manuscript. All authors edited and approved the final manuscript.

