## Supporting Information for "Intracellular phase separation of globular proteins facilitated by short cationic peptides"

### Materials and Methods

#### Cloning

The primers used to clone positive tags onto the C-terminus of GFP were purchased from Integrated DNA technologies (IDT). Primer sequences (denoted 5' to 3') and template plasmids used are shown below. Restriction enzyme sites were engineered into the primer sequence, with the forward primer containing a NcoI cut site and the reverse primers containing a XhoI cut site. All GFP variants contained a N-terminal 6xHis tag.

The following was used as the forward primer for all GFP variants: GATATACCATGGGTCATCACCACCACC

For the cloning of GFP variants, all reverse primers were designed to contain the following sites:
XhoI cut site STOP (Tag)Tag Plasmid Binding

pVY-1: sfGFP-tag1
GFP Template: sfGFP plasmid
Reverse Primer: TAATCTCGAGTTATCTTTTCTTCCTTTTCTTTGAACCGCCTTTCTTGTACAGCTCGTC-CA

pVY-2: sfGFP-tag2
GFP Template: sfGFP plasmid
Reverse Primer: TAATCTCGAGTTATCTTTTCTTCCTTTTCTTTGAACCGCCTCTTTTCTTCCTTTTCTTT-GAACCGCCTTTCTTGTACAGCTCGTCCATTC

pVY-3: sfGFP-tag3
GFP Template: pVY-1
Reverse Primer:
ATACTCGAGTTATTAACGCTTCTTACGCTTTTTGCTACCGCCACGCTTCTTACGCTTTTTGCTACCG-CCTCTTTTCTTCCTTTTCTTTGAA

pVY-4: GFP(0)-tag1
GFP Template: GFP(0) plasmid
Reverse Primer:
ATACTCGAGTTATTAACGCTTCTTACGCTTTTTGCTACCGCCCTTGTAGCGTTCGTCCATTC

pVY-5: GFP(0)-tag2
GFP Template: GFP(0) plasmid
Reverse Primer: ATACTCGAGCTATTAACGCTTCTTACGCTTTTTGCTACCGCCACGCTTCTTGC-GTTTCTTGCTTCCTCCCTTGTAGCGTTCGTCCATTC

pVY-6: GFP(0)-tag3
GFP template: pVY-5
Reverse Primer: ATACTCGAGTTATTATCGCTTTTTACGTTTCTTTGAACCTCCACGCTTCTTAC-GCTTTTTGCTACCGCCACGCTT

pVY-7: GFP(+6)-tag1
GFP template: GFP(+6) plasmid
Reverse Primer: ATACTCGAGTTATTAACGCTTCTTACGCTTTTTGCTACCGCCCTTGTAGCGTTCGTC-CATTC (same as pVY-4 reverse primer)

VY-8: GFP(+6)-tag2

GFP template: GFP(+6) plasmid

Reverse Primer: ATACTCGAGCTATTAACGCTTCTTACGCTTTTGCTACCGCCACGCTTCTTGC-GTTTCTTGCTTCCTCCCTTGTAGCGTTCGTCCATTC (same as pVY-5 reverse primer)

pVY-9: GFP(+6)-tag3

Template: pVY-8

Reverse Primer: ATACTCGAGTTATTATCGCTTTTACGTTTCTTTGAACCTCCACGCTTCTTAC-GCTTTTTGCTACCGCCACGCTT (same as pVY-6 reverse primer)

Primers were diluted to 10 μM in Milli-Q water. PCR reactions were performed using Phusion polymerase as instructed by New England Biolabs (NEB). PCR products were purified using the QIAquick PCR purification kit (Qiagen). Purified PCR products were digested with restriction enzymes, NcoI and XhoI. DpnI was added to the digestion of PCR amplified GFP inserts but not to the digestion of the vector. Reactions were run on an 1% agarose gel (TopVision agarose) and relevant bands were then excised and purified using a QIAquick gel extraction kit (Qiagen). The final plasmid was assembled using T4 ligase (NEB) using a 5:1 molar ratio of insert to vector. 2 μL of the ligation reaction was transformed into chemically competent NEB5α cells. Following sequence verification by Genewiz, plasmids were transformed into NiCo21(DE3) cells (NEB) for protein expression and *in vivo* assays.

For the following plasmids, mScarletI-tag was amplified from DNA sequences purchased from TWIST biosciences. Vector templates and amplified PCR products were then digested using restriction enzymes and loaded on a 1% agarose gel (TopVision agarose). The relevant bands were then excised and purified as described above. Plasmids were assembled using T4 ligase at a 3:1 molar ratio of insert to vector, transformed into NEB5α and NiCo21(DE3) cells, and sequence verified.

pVY-10: mScarletI-tag2

mScarletI-tag2 TWIST template: GAAGTGCCATTCCGCCTGACCTATAAGGCTCGTATGATATATTCAG-GGAGACCACAACGGTTTCCCTCTACAAATAATTTTGTTTAACTTTCTAGATTTAAGAAGGAGA-TATACATATGGGTCATCACCACCACCATCACGGTGGCGCTAGTAAAGGAGAAGCTGTGATTAAA-GAGTTCATGCGCTTCAAAGTTCACATGGAGGGTTCTATGAACGGTCACGAGTTCGAGATCGAAGG-CGAAGGCGAGGGCCGTCCGTATGAAGGCACCCAGACCGCCAAACTGAAAGTGACTAAAGGCGGC-CCGCTGCCTTTTTCCTGGGACATCCTGAGCCCGCAATTTATGTACGGTTCTAGGGCGTTCATCAAA-CACCCAGCGGATATCCCGGACTATTATAAGCAGTCTTTTCCGGAAGGTTTCAAGTGGGAACGCGTA-ATGAATTTTGAAGATGGTGGTGCCGTGACCGTCACTCAGGACACCTCCCTGGAGGATGGCACCCT-GATCTATAAAGTTAAACTGCGTGGTACTAATTTTCCACCTGATGGCCCGGTGATGCAGAAAAAGAC-GATGGGTTGGGAGGCGTCTACCGAACGCTTGTATCCGGAAGATGGTGTGCTGAAAGGCGACATTA-AAATGGCCCTGCGCCTGAAAGATGGCGGCCGCTATCTGGCTGACTTCAAAACCACGTACAAAG-CCAAGAAACCTGTGCAGATGCCTGGCGCGTACAATGTGGACCGCAAACTGGACATCACCTCT-CATAATGAAGATTATACGGTGGTAGAGCAATATGAGCGCTCCGAGGGTCGTCATTCTACCGGTGG-CATGGATGAACTATACAAAGGAGGAAGCAAGAAACGCAAGAAGCGTGGCGGTAGCAAAAAGCG-TAAGAAGCGTTAATAAACTTAATTAAGGATCCGAATTCGAGCTCCGTCGACAAGCTTGCGGCCG-CACTCGAGCACCTGCAGGCATGCAAGCTCTAGAGGCAAGGCTAGGTGGAGGCTCAGTG

Forward Primer: ATAAGGCTCGTATGATATATTCAGG

Reverse Primer: TGCCTCTAGAGCTTGCAT

Vector Template: pET-24a-mScarletI-v2 (plasmid map can be provided upon request)

Restriction Enzymes: NdeI and XhoI

pVY-11: mScarletI-tag3

mScarletI-tag3 TWIST template: GAAGTGCCATTCCGCCTGACCTATAAGGCTCGTATGATATATTCAG-GGAGACCACAACGGTTTCCCTCTACAAATAATTTTGTTTAACTTTCTAGATTTAAGAAGGAGA-TATACATATGGGTCATCACCACCACCATCACGGTGGCGCTAGTAAAGGAGAAGCTGTGATTAAA-GAGTTCATGCGCTTCAAAGTTCACATGGAGGGTTCTATGAACGGTCACGAGTTCGAGATCGAAGG-CGAAGGCGAGGGCCGTCCGTATGAAGGCACCCAGACCGCCAAACTGAAAGTGACTAAAGGCGGC-CCGCTGCCTTTTTCCTGGGACATCCTGAGCCCGCAATTTATGTACGGTTCTAGGGCGTTCATCAAA-CACCCAGCGGATATCCCGGACTATTATAAGCAGTCTTTTCCGGAAGGTTTCAAGTGGGAACGCGTA-ATGAATTTTGAAGATGGTGGTGCCGTGACCGTCACTCAGGACACCTCCCTGGAGGATGGCACCCT-GATCTATAAAGTTAAACTGCGTGGTACTAATTTTCCACCTGATGGCCCGGTGATGCAGAAAAAGAC-GATGGGTTGGGAGGCGTCTACCGAACGCTTGTATCCGGAAGATGGTGTGCTGAAAGGCGACATTA-AAATGGCCCTGCGCCTGAAAGATGGCGGCCGCTATCTGGCTGACTTCAAAACCACGTACAAAGC-CAAGAAACCTGTGCAGATGCCTGGCGCGTACAATGTGGACCGCAAACTGGACATCACCTCTCATA-ATGAAGATTATACGGTGGTAGAGCAATATGAGCGCTCCGAGGGTCGTCATTCTACCGGTGGCAT-GGATGAACTATACAAAGGAGGAAGCAAGAAACGCAAGAAGCGTGGCGGTAGCAAAAAGCGTA-AGAAGCGTGGAGGTTCAAAGAAACGTAAAAAGCGATAATAAACTTAATTAAGGATCCGAATTC-GAGCTCCGTCGACAAGCTTGCGGCCGCACTCGAGCACCTGCAGGCATGCAAGCTCTAGAGGCAAG-GCTAGGTGGAGGCTCAGTG

Forward Primer: ATAAGGCTCGTATGATATATTCAGG (same as pVY-10 forward primer)

Reverse Primer: TGCCTCTAGAGCTTGCAT (same as pVY-10 reverse primer)

Vector template: pET24a-mScarletI-v2 (plasmid map can be provided upon request)

Restriction Enzymes: NdeI and XhoI

pVY-12: CAT-tag3

CAT-tag3 TWIST template:
GAAGTGCCATTCCGCCTGACCTCTTTAAGAAGGAGATATACCATGGGTCATCACCACCACCATCAC-GGTGGCGCTGAGAAAAAAATCACTGGATATACCACCGTTGATATATCCCAATGGCATCGTAAAGAA-CATTTTGAGGCATTTCAGTCAGTTGCTCAATGTACCTATAACCAGACCGTTCAGCTGGATATTACGG-CCTTTTTAAAGACCGTAAAGAAAAATAAGCACAAGTTTTATCCGGCCTTTATTCACATTCTTGCCCG-CCTGATGAATGCTCATCCGGAATTCCGTATGGCAATGAAAGACGGTGAGCTGGTGATATGGGATAGT-GTTCACCCTTGTTACACCGTTTTCCATGAGCAAACTGAAACGTTTTCATCGCTCTGGAGTGAATAC-CACGACGATTTCCGGCAGTTTCTACACATATATTCGCAAGATGTGGCGTGTTACGGTGAAAACCTG-GCCTATTTCCCTAAAGGGTTTATTGAGAATATGTTTTTCGTCTCAGCCAATCCCTGGGTGAGTTTCAC-CAGTTTTGATTTAAACGTGGCCAATATGGACAACTTCTTCGCCCCCGTTTTCACCATGGGCAAATAT-TATACGCAAGGCGACAAGGTGCTGATGCCGCTGGCGATTCAGGTTCATCATGCCGTCTGTGATG-GCTTCCATGTCGGCAGAATGCTTAATGAATTACAACAGTACTGCGATGAGTGGCAGGGCGGGG-CGGGAGGAAGCAAGAAACGCAAGAAGCGTGGCGGTAGCAAAAAGCGTAAGAAGCGTGGAG-GTTCAAAGAAACGTAAAAAGCGATAATAACTCGAGTCTGGTAAAGAAACCGCTGCAGGCTAGGTG-GAGGCTCAGTG

Forward Primer: CTTTAAGAAGGAGATATACCATGGG

Reverse Primer: GCAGCGGTTTCTTTACCAGA

Vector Template: sfGFP plasmid

Restriction Enzymes: NcoI and XhoI

Note: for the cloning of CAT-tag3, only the vector template was digested with restriction enzymes. The digested vector was then mixed with purified CAT-tag3 insert obtained by PCR amplification of the TWIST template using the forward and reverse primers listed. The insert and backbone were annealed using NEBuilder HiFi DNA Assembly (described below).

The following plasmids were constructed using HiFi Assembly. Briefly, PCR reactions were performed and PCR products were purified as described above. Purified fragments were assembled using NEBuilder HiFi DNA Assembly Master Mix and were annealed at 50 °C for 15 min. The assembled plasmid was transformed into NEB5α and following sequence verification into NiCo21(DE3) cells. The sequences of plasmids were confirmed by Sanger sequencing (Genewiz).

pVY-13: mScarletI
mScarletI Template: pEB2-mScarletI from Addgene (#104007)
Forward Primer: TTTAAGAAGGAGGAGTACATATGGGTCATCACCACCACCATCACGGTGGCGCTAG-TAAAGGAGAAGCTGTGATTAAAG
Reverse Primer: CTTTGTTAGCAGCCGGATCTTTATTATTTGTATAGTTCATCCATGC
pET24a vector Template: pET24a-ELP[V-30] from Addgene (#67014)
Forward Primer: AGCGCCACCGTGATGGTGGTGGTGATGACCCATATGTACTCCTCCTTCTTAAAGTTA-AACAAAAT
Reverse Primer: TAAAGATCCGGCTGCTAACAAAGC

pVY-14: TC-CAT
CAT Template: CAT plasmid[1]
Forward Primer: ACTTTAAGAAGGAGATATACATGGGTTGTTGCCCTGGGTGTTGTGGTGGCGCT-GAGAAAAAAATCACTGGATATAC
Reverse Primer: ATGAGTATTCAACATTTCCG
pET24a vector template: CAT plasmid
Vector Forward Primer: CGGAAATGTTGAATACTCATACTCTTCCTTTTTCAATCATG
Vector Reverse Primer: GTATATCTCCTTCTTAAAGTTAAAC

pVY-15: TC-CAT-tag3
CAT Template: pVY-12
Forward Primer: ACTTTAAGAAGGAGATATACATGGGTTGTTGCCCTGGGTGTTGTGGTGGCGCT-GAGAAAAAAATCACTGGATATAC
Reverse Primer: ATGAGTATTCAACATTTCCG
pET24a vector template: CAT plasmid
Vector Forward Primer: CGGAAATGTTGAATACTCATACTCTTCCTTTTTCAATCATG (same as pVY-14 vector forward primer)
Vector Reverse Primer: GTATATCTCCTTCTTAAAGTTAAAC (same as pVY-14 vector reverse primer)

pVY-16: TC-TEM1
Template: sfGFP plasmid
Forward Primer: TTTAAGAAGGAGATATACATATGGGTTGTTGCCCTGGGTGTTGTGGTGGCGCTAG-TATTCAACATTTCCGTG
Reverse Primer: GCCGTTAATTAAGTTTATTACCAATGCTTAATCAGTGAG
pBAD vector template: p2-39 mScarletI-tag2 (plasmid map can be provided upon request)
Vector Forward Primer: TAATAAACTTAATTAACGGCAC
Vector Reverse Primer: ATGTATATCTCCTTCTTAAAGTTAAAC

pVY-17: TC-TEM1-tag3
Template: sfGFP plasmid
Forward Primer: TTTAAGAAGGAGATATACATATGGGTTGTTGCCCTGGGTGTTGTGGTGGCGCTAG-TATTCAACATTTCCGTG
Reverse Primer: TTGCGTTTCTTGCTTCCTCCCCAATGCTTAATCAGTGAG
pBAD vector template: p2-39 mScarletI-tag2 (plasmid map can be provided upon request)
Vector Forward Primer: GGAGGAAGCAAGAAACGC
Vector Reverse Primer: ATGTATATCTCCTTCTTAAAGTTAAACAAAATTATTTC

pVY-18: TC-sfGFP
TC-sfGFP Template: sfGFP plasmid
Forward Primer: ACTTTAAGAAGGAGATATACCATGGGTTGTTGCCCTGGGTGTTGTGGTGGCGCTAG-CAAAGGTGAAGAG
Reverse Primer: ATGAGTATTCAACATTTCCG
pET vector template: sfGFP plasmid
Vector Forward Primer: CGGAAATGTTGAATACTCATACTCTTCCTTTTTCAATCATG
Vector Reverse Primer: GTATATCTCCTTCTTAAAGTTAAAC

pVY-19: TC-sfGFP-tag3
TC-sfGFP-tag3 Template: pVY-3
Forward Primer: ACTTTAAGAAGGAGATATACCATGGGTTGTTGCCCTGGGTGTTGTGGTGGCGCTAG-CAAAGGTGAAGAG (same as pVY-18 forward primer)
Reverse Primer: ATGAGTATTCAACATTTCCG (same as pVY-18 reverse primer)
pET vector template: pVY-3
Vector Forward Primer: CGGAAATGTTGAATACTCATACTCTTCCTTTTTCAATCATG (same as pVY-18 vector forward primer)
Vector Reverse Primer: GTATATCTCCTTCTTAAAGTTAAAC (same as pVY-18 vector reverse primer)

Amino Acid Sequence of GFP, mScarletI, TEM1, and CAT variants:
His6x and TC tags with linkers (GGA) are underlined. Cationic tags are **bolded and underlined.** Tagged sfG-FP variants contain an additional lysine residue (**bolded** but not underlined) between the C-terminus of sfGFP and N-terminus of the peptide tag because sfGFP has a net charge of -7. This lysine residue was included to ensure charge-equivalence between respective tagged variants and was not added to variants containing GFP(0) or GFP(+6) globular domains.

His6x-sfGFP-tag1
MGHHHHHHGGASKGEELFTGVVPILVELDGDVNGHKFSVRGEGEGDATNGKLTLKFICTTGKLPVP-WPTLVTTLTYGVQCFSRYPDHMKQHDFFKSAMPEGYVQERTISFKDDGTYKTRAEVKFEGDTLVNRI-ELKGIDFKEDGNILGHKLEYNFNSHNVYITADKQKNGIKANFKIRHNVEDGSVQLADHYQQNTPIGDG-PVLLPDNHYLSTQSALSKDPNEKRDHMVLLEFVTAAGITHGMDELYK**KGGSKKRKKR**

His6x-sfGFP-tag2
MGHHHHHHGGASKGEELFTGVVPILVELDGDVNGHKFSVRGEGEGDATNGKLTLKFICTTGKLPVP-WPTLVTTLTYGVQCFSRYPDHMKQHDFFKSAMPEGYVQERTISFKDDGTYKTRAEVKFEGDTLVNRI-ELKGIDFKEDGNILGHKLEYNFNSHNVYITADKQKNGIKANFKIRHNVEDGSVQLADHYQQNTPIGDG-PVLLPDNHYLSTQSALSKDPNEKRDHMVLLEFVTAAGITHGMDELYK**KGGSKKRKKRGGSKKRK-KR**

His6x-sfGFP-tag3
MGHHHHHHGGASKGEELFTGVVPILVELDGDVNGHKFSVRGEGEGDATNGKLTLKFICTTGKLPVP-WPTLVTTLTYGVQCFSRYPDHMKQHDFFKSAMPEGYVQERTISFKDDGTYKTRAEVKFEGDTLVNRI-ELKGIDFKEDGNILGHKLEYNFNSHNVYITADKQKNGIKANFKIRHNVEDGSVQLADHYQQNTPIGDG-PVLLPDNHYLSTQSALSKDPNEKRDHMVLLEFVTAAGITHGMDELYK**KGGSKKRKKRGGSKKRK-KRGGSKKRKKR**

His6x-GFP(0)-tag1
MGHHHHHHGGASKGERLFTGVVPILVELDGDVNGHKFSVRGEGEGDATNGKLTLKFICTTGKLPVP-WPTLVTTLTYGVQCFSRYPDHMKQHDFFKSAMPEGYVQERTISFKDDGTYKTRAEVKFEGDTLVNRI-ELKGRDFKEDGNILGHKLEYNFNSHNVYITADKQKNGIKANFKIRHNVEDGSVQLADHYQQNTPIG-

DGPVLLPRNHYLSTQSALSKDPKEKRDHMVLLEFVTAAGITHGMDERYK**GGSKKRKKR**

His6x-GFP(0)-tag2
MGHHHHHHGGASKGERLFTGVVPILVELDGDVNGHKFSVRGEGEGDATNGKLTLKFICTTGKLPVP-WPTLVTTLTYGVQCFSRYPDHMKQHDFFKSAMPEGYVQERTISFKDDGTYKTRAEVKFEGDTLVNRI-ELKGRDFKEDGNILGHKLEYNFNSHNVYITADKQKNGIKANFKIRHNVEDGSVQLADHYQQNTPIG-DGPVLLPRNHYLSTQSALSKDPKEKRDHMVLLEFVTAAGITHGMDERYK**GGSKKRKKRGGSKKRK-KR**

His6x-GFP(0)-tag3
MGHHHHHHGGASKGERLFTGVVPILVELDGDVNGHKFSVRGEGEGDATNGKLTLKFICTTGKLPVP-WPTLVTTLTYGVQCFSRYPDHMKQHDFFKSAMPEGYVQERTISFKDDGTYKTRAEVKFEGDTLVNRI-ELKGRDFKEDGNILGHKLEYNFNSHNVYITADKQKNGIKANFKIRHNVEDGSVQLADHYQQNTPIG-DGPVLLPRNHYLSTQSALSKDPKEKRDHMVLLEFVTAAGITHGMDERYK**GGSKKRKKRGGSKKRK-KRGGSKKRKKR**

His6x-GFP(+6)-tag1
MGHHHHHHGGASKGERLFTGVVPILVELDGDVNGHKFSVRGEGEGDATNGKLTLKFICTTGKLPVP-WPTLVTTLTYGVQCFSRYPKHMKQHDFFKSAMPEGYVQERTISFKKDGTYKTRAEVKFEGRTLVNRI-ELKGRDFKEDGNILGHKLEYNFNSHNVYITADKQKNGIKANFKIRHNVEDGSVQLADHYQQNTPIG-DGPVLLPRNHYLSTQSALSKDPKEKRDHMVLLEFVTAAGITHGMDERYK**GGSKKRKKR**

His6x-GFP(+6)-tag2
MGHHHHHHGGASKGERLFTGVVPILVELDGDVNGHKFSVRGEGEGDATNGKLTLKFICTTGKLPVP-WPTLVTTLTYGVQCFSRYPKHMKQHDFFKSAMPEGYVQERTISFKKDGTYKTRAEVKFEGRTLVNRI-ELKGRDFKEDGNILGHKLEYNFNSHNVYITADKQKNGIKANFKIRHNVEDGSVQLADHYQQNTPIG-DGPVLLPRNHYLSTQSALSKDPKEKRDHMVLLEFVTAAGITHGMDERYK**GGSKKRKKRGGSKKRK-KR**

His6x-GFP(+6)-tag2
MGHHHHHHGGASKGERLFTGVVPILVELDGDVNGHKFSVRGEGEGDATNGKLTLKFICTTGKLPVP-WPTLVTTLTYGVQCFSRYPKHMKQHDFFKSAMPEGYVQERTISFKKDGTYKTRAEVKFEGRTLVNRI-ELKGRDFKEDGNILGHKLEYNFNSHNVYITADKQKNGIKANFKIRHNVEDGSVQLADHYQQNTPIG-DGPVLLPRNHYLSTQSALSKDPKEKRDHMVLLEFVTAAGITHGMDERYK**GGSKKRKKRGGSKKRK-KRGGSKKRKKR**

His6x-mScarletI
MGHHHHHHGGASKGEAVIKEFMRFKVHMEGSMNGHEFEIEGEGEGRPYEGTQTAKLKVTKGG-PLPFSWDILSPQFMYGSRAFIKHPADIPDYYKQSFPEGFKWERVMNFEDGGAVTVTQDTSLEDGTLI-YKVKLRGTNFPPDGPVMQKKTMGWEASTERLYPEDGVLKGDIKMALRLKDGGRYLADFKTTYKAK-KPVQMPGAYNVDRKLDITSHNEDYTVVEQYERSEGRHSTGGMDELYK

His6x-mScarletI-tag2
MGHHHHHHGGASKGEAVIKEFMRFKVHMEGSMNGHEFEIEGEGEGRPYEGTQTAKLKVTKGG-PLPFSWDILSPQFMYGSRAFIKHPADIPDYYKQSFPEGFKWERVMNFEDGGAVTVTQDTSLEDGTLI-YKVKLRGTNFPPDGPVMQKKTMGWEASTERLYPEDGVLKGDIKMALRLKDGGRYLADFKTTYKAK-KPVQMPGAYNVDRKLDITSHNEDYTVVEQYERSEGRHSTGGMDELYK**GGSKKRKKRGGSKKRKKR**

His6x-mScarletI-tag3
MGHHHHHHGGASKGEAVIKEFMRFKVHMEGSMNGHEFEIEGEGEGRPYEGTQTAKLKVTKGG-PLPFSWDILSPQFMYGSRAFIKHPADIPDYYKQSFPEGFKWERVMNFEDGGAVTVTQDTSLEDGTLI-YKVKLRGTNFPPDGPVMQKKTMGWEASTERLYPEDGVLKGDIKMALRLKDGGRYLADFKTTYKA-KKPVQMPGAYNVDRKLDITSHNEDYTVVEQYERSEGRHSTGGMDELYK**GGSKKRKKRGGSKKRK-KRGGSKKRKKR**

TC-CAT
MGCCPGCCGGAEKKITGYTTVDISQWHRKEHFEAFQSVAQCTYNQTVQLDITAFLKTVKKNKHK-FYPAFIHILARLMNAHPEFRMAMKDGELVIWDSVHPCYTVFHEQTETFSSLWSEYHDDFRQFLHI-YSQDVACYGENLAYFPKGFIENMFFVSANPWVSFTSFDLNVANMDNFFAPVFTMGKYYTQGDKVLM-PLAIQVHHAVCDGFHVGRMLNELQQYCDEWQGGA

TC-CAT-tag3
MGCCPGCCGGAEKKITGYTTVDISQWHRKEHFEAFQSVAQCTYNQTVQLDITAFLKTVKKNKHK-FYPAFIHILARLMNAHPEFRMAMKDGELVIWDSVHPCYTVFHEQTETFSSLWSEYHDDFRQFLHI-YSQDVACYGENLAYFPKGFIENMFFVSANPWVSFTSFDLNVANMDNFFAPVFTMGKYYTQGDKVLM-PLAIQVHHAVCDGFHVGRMLNELQQYCDEWQGGA**GGSKKRKKRGGSKKRKKRGGSKKRKKR**

TC-TEM1
MGCCPGCCGGASIQHFRVALIPFFAAFCLPVFAHPETLVKVKDAEDQLGARVGYIELDLNSGKILES-FRPEERFPMMSTFKVLLCGAVLSRIDAGQEQLGRRIHYSQNDLVEYSPVTEKHLTDGMTVRELCSAA-ITMSDNTAANLLLTTIGGPKELTAFLHNMGDHVTRLDRWEPELNEAIPNDERDTTMPVAMATTL-RKLLTGELLTLASRQQLIDWMEADKVAGPLLRSALPAGWFIADKSGAGERGSRGIIAALGPDGKPSRIV-VIYTTGSQATMDERNRQIAEIGASLIKHW

TC-TEM1-tag3
MGCCPGCCGGASIQHFRVALIPFFAAFCLPVFAHPETLVKVKDAEDQLGARVGYIELDLNSGKILES-FRPEERFPMMSTFKVLLCGAVLSRIDAGQEQLGRRIHYSQNDLVEYSPVTEKHLTDGMTVRELCSAA-ITMSDNTAANLLLTTIGGPKELTAFLHNMGDHVTRLDRWEPELNEAIPNDERDTTMPVAMATTL-RKLLTGELLTLASRQQLIDWMEADKVAGPLLRSALPAGWFIADKSGAGERGSRGIIAALGPDGKPSRIV-VIYTTGSQATMDERNRQIAEIGASLIKHW**GGSKKRKKRGGSKKRKKRGGSKKRKKR**

TC-sfGFP
MGCCPGCCGGASKGEELFTGVVPILVELDGDVNGHKFSVRGEGEGDATNGKLTLKFICTTGKLPVP-WPTLVTTLTYGVQCFSRYPDHMKQHDFFKSAMPEGYVQERTISFKDDGTYKTRAEVKFEGDTLVNRI-ELKGIDFKEDGNILGHKLEYNFNSHNVYITADKQKNGIKANFKIRHNVEDGSVQLADHYQQNTPIGDG-PVLLPDNHYLSTQSALSKDPNEKRDHMVLLEFVTAAGITHGMDELYK

TC-sfGFP-tag3
MGCCPGCCGGASKGEELFTGVVPILVELDGDVNGHKFSVRGEGEGDATNGKLTLKFICTTGKLPVP-WPTLVTTLTYGVQCFSRYPDHMKQHDFFKSAMPEGYVQERTISFKDDGTYKTRAEVKFEGDTLVNRI-ELKGIDFKEDGNILGHKLEYNFNSHNVYITADKQKNGIKANFKIRHNVEDGSVQLADHYQQNTPIG-DGPVLLPDNHYLSTQSALSKDPNEKRDHMVLLEFVTAAGITHGMDELYK**GGSKKRKKRGGSKKRK-KRGGSKKRKKR**

| Tags | Tag Sequence | Net Charge |
|---|---|---|
| tag1 | GGSKKRKKR | +6 |
| tag2 | GGSKKRKKR GGSKKRKKR | +12 |
| tag3 | GGSKKRKKR GGSKKRKKR GGSKKRKKR | +18 |

**Electrostatic maps of GFP variants**

The PDB file for sfGFP (PDB ID: 2B3P) was obtained from the RCSB protein data bank and loaded in PyMOL v2.4.1 (Schrödinger, LLC). Using the PDB file for sfGFP as a template, GFP(0) and GFP(+6) were made by introducing residue substitutions in PyMOL. PDBs of the minimized peptide tag structures were obtained using PEP-FOLD3[2–4]that allows both (i. The C-terminus of the globular domain was appended to the N-terminus of the tags in PyMOL and electrostatic surface maps were generated using the ABPS Electrostatics plugin.

**Constraints for image analysis using MicrobeJ**

Briefly, images stacks were subjected to a rolling ball background subtraction with a 100 pixel radius. Stacks of images were individually thresholded using the Li method in MicrobeJ (version 15.3l (14)) and cells that met the area, length, width, and angularity constraints listed in the table below were identified and considered for analysis. The following constraints were used for all image stacks: circularity = 0.25 – max; curvature = 0 – max; sinuosity = 0 – max; solidity = 0 – max; intensity = 0 – max. Bacteria were detected using the fit shape, rod-shaped mode.

| Images | Area (pixels$^2$) | Length (pixels) | Width (pixels) | Angularity |
|---|---|---|---|---|
| Isotropic GFP variants at 2 and 8 h (unless specified otherwise) | 100-800 | 20 - max | 0-20 | 0 – 0.4 |
| Isotropic GFP variants at 24 h (unless specified otherwise) Also used for Tagged GFP variants with cell clumps at 2, 4, 6, and 24 h | 100-800 | 20 - max | 0-15 | 0 – 0.4 |
| Isotropic GFP(+24) and (+36) at 8 and 24 h | 100-800 | 25 - max | 0-20 | 0 – 0.4 |
| Tagged GFP variants at 2, 4, 6, 24 h (unless otherwise specified) | 100-800 | 10 - max | 0-25 | 0 – 0.45 |
| mScarletI variants at 2, 4, 24 h | 100-2000 | 25 - max | 0-25 | 0 – 0.45 |

**Image analysis using a custom MATLAB script**

Briefly, the nine central pixels of each cell image were averaged to determine the approximate average pixel intensity for non-condensate areas for each cell. All pixels in a cell with 20% higher intensity than this approximated average intensity were classified as condensates. This intensity threshold was selected after testing intensity thresholds from 0 to 30% higher than the approximated average intensity on isotropic GFP variants that had been separately categorized as condensate-containing or non-condensate containing[5]. The 20% threshold most accurately and clearly distinguished condensate-containing variants from non-condensate-containing variants.  A duplicate image with all non-condensate pixels set to zero was then created for each cell. The number of non-zero pixels in the duplicate image was divided by the number of pixels in the original image to determine the percentage of each cell that contained condensates. This percentage was then used to determine the number of condensate-containing cells by counting cells with a non-zero percentage as condensate-containing.. The number of condensate-containing cells was thus counted for each experiment and divided by the total number of cells identified by MicrobeJ to determine the fraction of condensate-containing cells.

**Cell growth assay**

Cells were grown as described in the "Protein expression" section and aliquots were collected at 0, 2, 4, 6, 8, 10, 12, 20, 22, and 24 h after induction. At each time point, aliquots of each sample were diluted 5- or 10-fold with LB media to a total volume of 1 mL. 150 μL of each diluted sample was then transferred to a 96-well black flat bottom tissue culture-treated plate (Corning REF#3904) in triplicate. GFP fluorescence intensity and Tecan $OD_{600}$ were measured for all samples in a plate reader (Tecan Infinite M200 Pro; excitation at 488 nm, emission at 530 nm, gain: 75). For mScarletI variants, Tecan $OD_{750}$ and RFP fluorescence intensity were measured (excitation at 569 nm, emission at 593 nm, gain: 75). The fluorescence intensity and Tecan OD of wells containing LB media were subtracted from raw fluorescence intensity values and raw Tecan OD values, respectively. Background subtracted fluorescence intensity values were then divided by the adjusted Tecan OD. This ratio of fluorescence intensity to Tecan OD was averaged across three biological replicates for each protein variant and then normalized to the highest average fluorescence intensity/Tecan OD ratio amongst all GFP variants (GFP(0), 22 h).

**Solubility assay**

Cell cultures (1 L) were grown and expressed as described in the "Protein Expression" section above. Cultures were incubated at 25 ºC for 24 h after induction with IPTG. 900 mL of each culture was then harvested by centrifugation (Thermo Scientific Sorvall Legend XTR) at 4000 rpm for 10 min in a swinging bucket rotor (Thermo Scientific TX-750). The cell pellet was resuspended in 13.5 mL of cell lysis buffer without NaCl (50 mM $NaH_2PO_4$, pH 8.0) and divided into 4.5 mL aliquots. All samples were subjected to one freeze-thaw cycle before being lysed by sonication on ice (2 s on, 4 s off at 40% amplitude for 5 min of sonics) with a 1/8 in. (diameter) microtip probe. The lysate was then centrifuged at 10,000 rpm for 30 minutes at 4 °C in a fixed angle rotor (Thermo Scientific, Fiberlite F15-8x50cy). The pellets were washed by resuspending in 4.5 mL of lysis buffer without NaCl, divided into three 1.5 mL aliquots in 2 mL microcentrifuge tubes. Samples were then pelleted at 16,000 x g in a microcentrifuge (Thermo Scientific Sorvall Legend Micro 21R) for 30 minutes at 4 °C. The supernatant was removed, and 1.4 mL of cell lysis buffer containing different concentrations of NaCl (0 mM, 100 mM or 1 M) was then added to each of the pellets to solubilize the protein condensates. Finally, samples were centrifuged again at 16,000 x g for 30 minutes at 4 °C. All supernatants (soluble fractions) and pellets (insoluble fractions) were run on Bolt™ 4-12% Bis-Tris Plus gels (Invitrogen) and imaged on a Gel Doc XR+ System (Bio-Rad) to determine condensate solubility. Analysis of gel images was done as described previously[5]. Briefly, a 100-pixel radius rolling ball background subtraction was performed in Fiji for each gel image. Rectangular ROIs were drawn around the band corresponding to GFP or spanning the height of the entire lane to capture all proteins. Histograms were generated from these ROIs, and line segments were drawn at the base of the peaks. The area under the curve was measured in order to measure the band intensities of GFP and total protein for each lane. The fraction of GFP in the soluble fraction was determined by dividing the band intensity of GFP in the soluble fraction by the band intensity of total protein in the soluble fraction. For each variant, the fraction of GFP in the soluble fraction was then normalized in GraphPad Prism (version 9.1.0) to the fraction extracted in 1 M NaCl and plotted as the normalized ratio of GFP to total protein in the supernatant.

**Protein purification for *in vitro* tests**

Cell cultures (1 L) were grown and expressed as described in the "Protein Expression" section above. Cultures were incubated at 25 ºC for 16-18 h after induction with IPTG. Cultures were then harvested by centrifugation (Thermo Scientific Sorvall Legend XTR) at 4000 rpm for 10 min in a swinging bucket rotor (Thermo Scientific TX-750). The cell pellet was resuspended in lysis buffer (50 mM $NaH_2PO_4$ , 300 mM NaCl, pH 8.0) and lysed by sonication on ice (2 s on, 4 s off at 60% amplitude for 8 min of sonics) using a 0.5 inch probe. The lysate was then centrifuged at 10,000 rpm for 30 min and the supernatant (clarified lysate) was collected for purification using immobilized metal affinity chromatography. 8 mL of His-Pur Ni-NTA slurry (Thermo Scientific) was used to purify protein harvested from 1 L cell culture. Briefly, the resin was equilibrated with lysis buffer and the clarified lysate was incubated with the resin for 5-10 min at room temperature. The resin was then washed in lysis buffer containing 50 mM imidazole and His-tagged proteins were eluted with lysis buffer containing 250 mM imidazole. All flow through, wash, and elution fractions were collected for analysis on a Bolt™ 4-12% Bis-Tris Plus gel (Invitrogen). Pure fractions were concentrated using Amicon Ultra centrifugal filter units with a 10 kDa molecular weight cutoff (Millipore Sigma) and then dialyzed against a physiological buffer (70 mM $K_2HPO_4$, 60 mM KCl, 40 mM NaCl, pH 7.4), which mimics the cytoplasmic ion concentrations in *E. coli.* The buffer was changed 7 times with at least 3 h equilibration between each buffer exchange.

**MALDI-TOF mass spectrometry**

The mass of purified GFP(+6)-tag3 used for *in vitro* assays was verified by MALDI-TOF mass spectrometry prior to use in experiments. As reported previously, salts were removed from the protein stock solution by performing a buffer exchange into Milli-Q water using 10 kDa MWCO Amicon Ultra centrifugal filter (Millipore Sigma)[5]. A 10 mg/mL sinapinic acid matrix (in 7:3 water:acetonitrile with 0.1% trifluoroacetic acid) was prepared and the GF-P(+6)-tag3 sample was premixed at a 4:6 sample:matrix ratio before being spotted on a grounded stainless steel target. A calibration was performed with Protein Standard II (Bruker) before collecting MALDI-TOF spectra on a Bruker ultrafleXtreme MALDI TOF/TOF. All spectra were obtained from the Columbia University Chemistry Department Mass Spectrometry Facility.

#### *In vitro* turbidity assays and microscopy of coacervates

Protein stock solutions were prepared at 1 mg/mL in physiological buffer (70 mM $K_2HPO_4$, 60 mM KCl, 40 mM NaCl, pH 7.4). 1 mg/mL RNA stock solutions were prepared fresh by dissolving total RNA from torula yeast type VI (Sigma-Aldrich) in physiological buffer and adjusting to pH 7.4. Both stock solutions were filtered using a 0.22 µm SFCA Thermo Scientific Nalgene 25 mm Syringe Filter. Protein and RNA stock solutions were mixed at varying mass ratios in a tissue culture-treated polystyrene 96-well half-area plate (Corning, REF#3697). Mixing ratios were prepared in triplicate, and absorbance at 600 nm was measured for each well on a plate reader (Tecan Infinite M200 Pro). Turbidity was then calculated using the equation, Turbidity = 100 - $10^{2-A}$. Following these measurements, GFP/RNA mixtures with high turbidity were prepared in a 384-well glass-bottom plate (Cellvis) and imaged on an Evos FL Auto 2 inverted fluorescence microscope (Invitrogen). Images were acquired using a 20X 0.40 NA Plan Fluor objective on GFP ( = 479-522 nm; = 525-550 nm; EVOS GFP light cube) and brightfield channels.

#### NaCl titration to determine critical salt concentration

Protein and RNA stock solutions were prepared in physiological buffer (70 mM $K_2HPO_4$, 60 mM KCl, 40 mM NaCl, pH 7.4) as described above. GFP(+6)-tag3 and RNA were mixed at the mass fraction that provided the highest turbidity, which was determined empirically (0.6 GFP mass fraction). The sample was prepared to a final volume of 1 mL in a 4 mL PMMA cuvette and stirred continuously at 22 °C, 500 rpm. A solution of 5 M NaCl dissolved in physiological buffer was titrated into the sample via 1 µL additions and the absorbance at 600 nm was collected following each addition using a Cary Ultraviolet/visible spectrophotometer. Measurements were collected until the absorbance stabilized. Absorbance was then converted to turbidity as described above and then plotted as a function of ionic strength in Prism GraphPad (version 9.1.0). Ionic strength was calculated using the formula: , where *n* is the number of ions in solution, $c_i$ is the concentration of ion species *i*, and $z_i$ is the valence of ion species *i*. For ionic strength calculations, we assumed the following dissociation reactions at pH 7.4: $K_2HPO_{4(s)} \rightarrow 2K^+_{(aq)} + HPO_4{}^{2-}{}_{(aq)}$; $KCl_{(s)} \rightarrow K^+_{(aq)} + Cl^-_{(aq)}$; and $NaCl_{(s)} \rightarrow Na^+_{(aq)} + Cl^-_{(aq)}$.

#### FlAsH staining of TC-AmpR and -AmpR-tag3 cultures

Overnight cultures were back diluted to $OD_{600}$ = 0.1 in 1 mL of LB media supplemented with 25 µg/mL chloramphenicol, 0.2% w/v arabinose, and 10 µM FlAsH-$EDT_2$ (Cayman Chemical) and incubated in 14 mL snap cap tubes in an incubator at 37 ºC with shaking at 225 rpm. After 2 h, the cultures were transferred to a 25 ºC incubator and allowed to shake at 225 rpm overnight. After 24 h, 100 µL of each culture was washed three times in an equal volume of BAL (British anti-Lewisite from Fisher Scientific) diluted to 1X in phosphate buffer saline, and adjusted to pH 7.4. Cells were washed in a microcentrifuge tube with centrifugation at 1,400 x g to collect the cell pellet between each washing step. Stained cells were then imaged on a 1% agarose pad as described in the "Optical Microscopy" section.

#### ReAsH-EDT staining of TC-CAT, TC-CAT-tag3, TC-sfGFP, and TC-sfGFP-tag3 cultures

Overnight cultures were back diluted to $OD_{600}$ ~ 0.1 in 2 mL of LB media supplemented with 100 µg/ml ampicillin, and incubated in 14 mL snap cap tubes in an incubator at 37 ºC with shaking at 225 rpm. After 2 h, the cultures reached $OD_{600}$ = 0.8 -1 and were induced with 1 mM IPTG. Cultures were subsequently stained with 1 µM ReAsH-$EDT_2$ (Invitrogen) and transferred to a 25 ºC incubator with shaking at 225 rpm overnight. 24 h after IPTG induction, 100 µL of each sample was collected in a microcentrifuge tube and washed three times in an equal volume of 1X BAL in phosphate buffer saline, pH 7.4 before imaging on a 1.5% LB agarose pad using the protocol described in the "Optical microscopy" section.

**Carbenicillin Sensitivity Spotting Assay**

Overnight cultures of TC-TEM1 and TC-TEM1-tag3 were prepared as described above and then diluted to $OD_{600}$ = 0.1 in LB media supplemented with 25 µg/ml chloramphenicol, 0.2 % (w/v) arabinose, and 10 µM FlAsH-$EDT_2$ (Fisher Scientific). Induced cultures were grown at 37 ºC, 225 rpm for 2 h. All cultures were normalized to the $OD_{600}$ of the least dense sample (~0.7) in LB media. Normalized cultures were serially diluted 20-fold in LB media. 5 µL of all normalized cultures and their dilutions (20X and 400X) were then spotted onto LB agar plates supplemented with either 0.2 % (w/v) arabinose or 10 mM glucose, 25 µg/mL chloramphenicol, and varying concentrations of carbenicillin (0, 25, 50, 100, or 200 µg/mL). Samples on plates were allowed to dry fully before incubation in the dark at 25 ºC for ~30 hours and then imaged on a Gel Doc XR+ System (Bio-Rad). Samples from the plate were also picked into 20 µL LB media and imaged on a microscope slide to confirm the presence of condensates (as described in "Optical Microscopy").

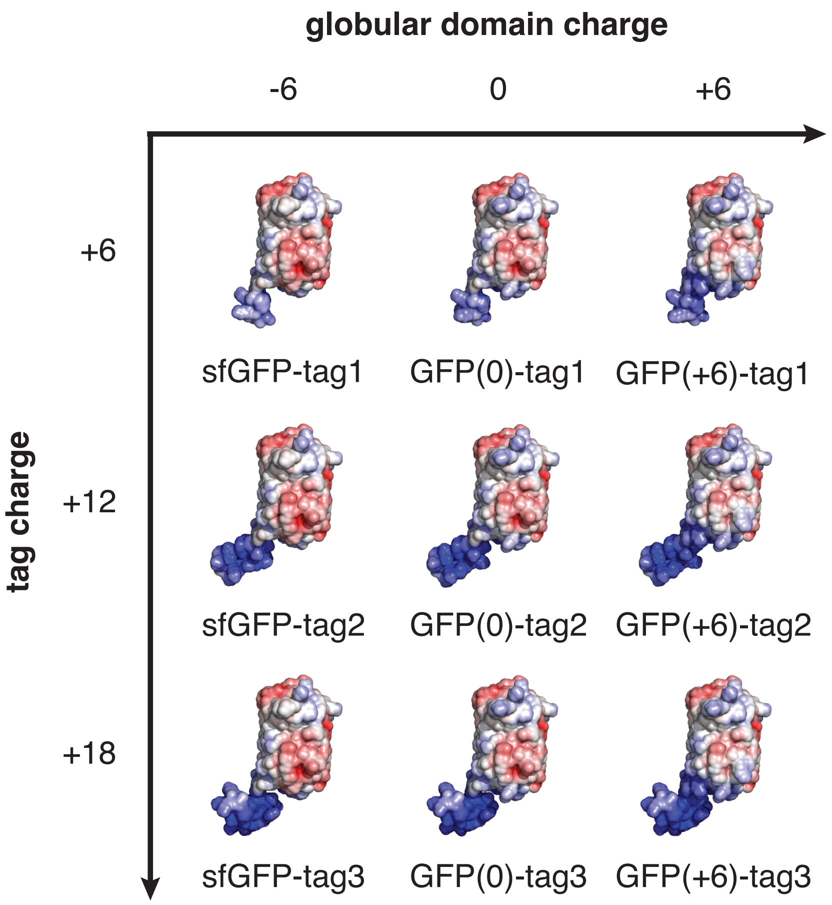

**Supplementary Figure 1. Electrostatic surface representations of the cationic GFP variants investigated in this study.** Each GFP variant is a combination of a globular domain (sfGFP, GFP(0), or GFP(+6)) and a cationic peptide tag, which consists of one or more repeats of the amino acid sequence, GGSKKRKKR. Additional sequence information can be found in the materials and methods. Electrostatic maps were generated using the ABPS Electrostatics plugin in PyMol.

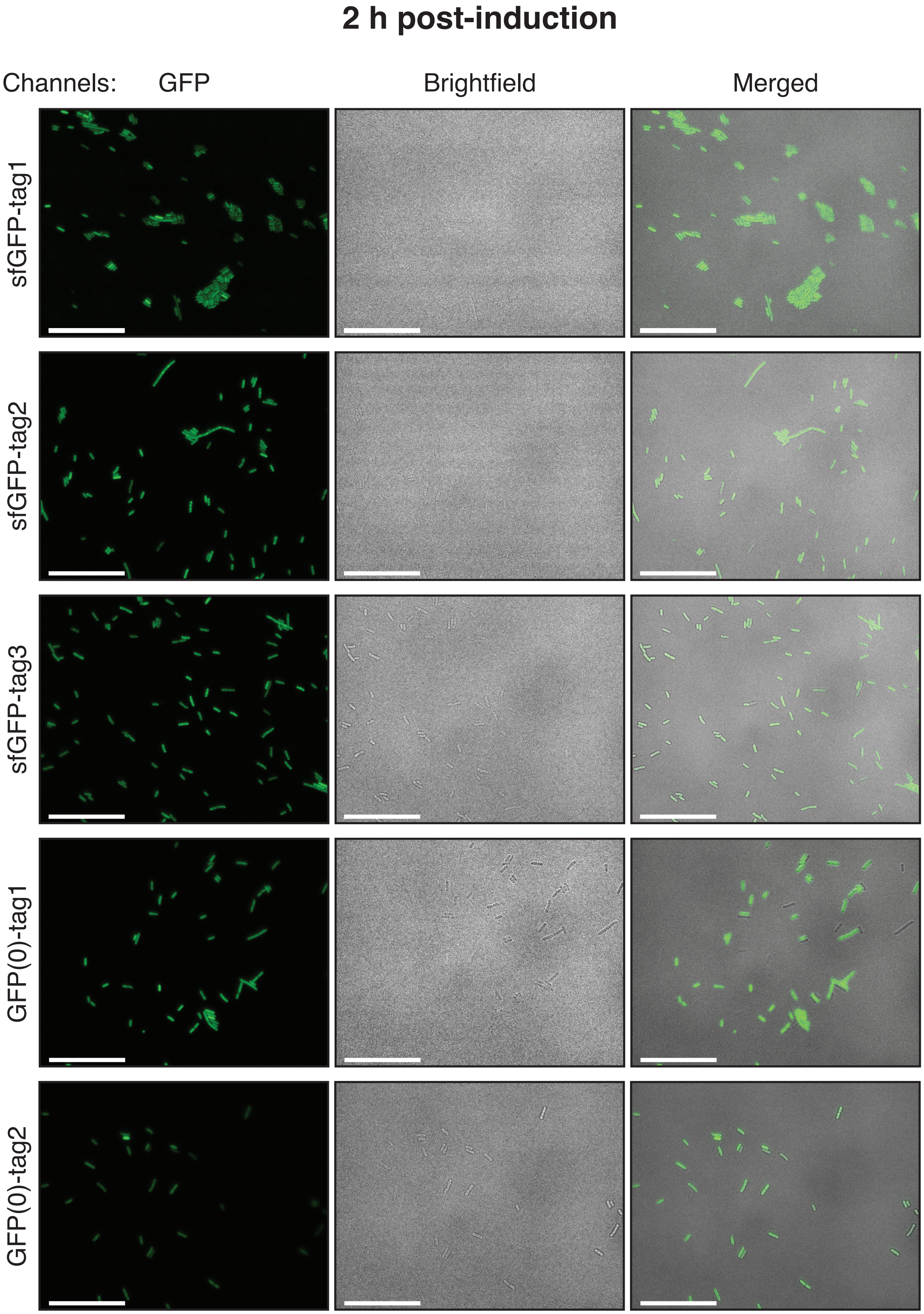

2 h post-induction
Channels:
GFP
Brightfield
Merged
sfGFP-tag1
sfGFP-tag2
sfGFP-tag3
GFP(0)-tag1
GFP(0)-tag2

### 2 h post-induction (continued)

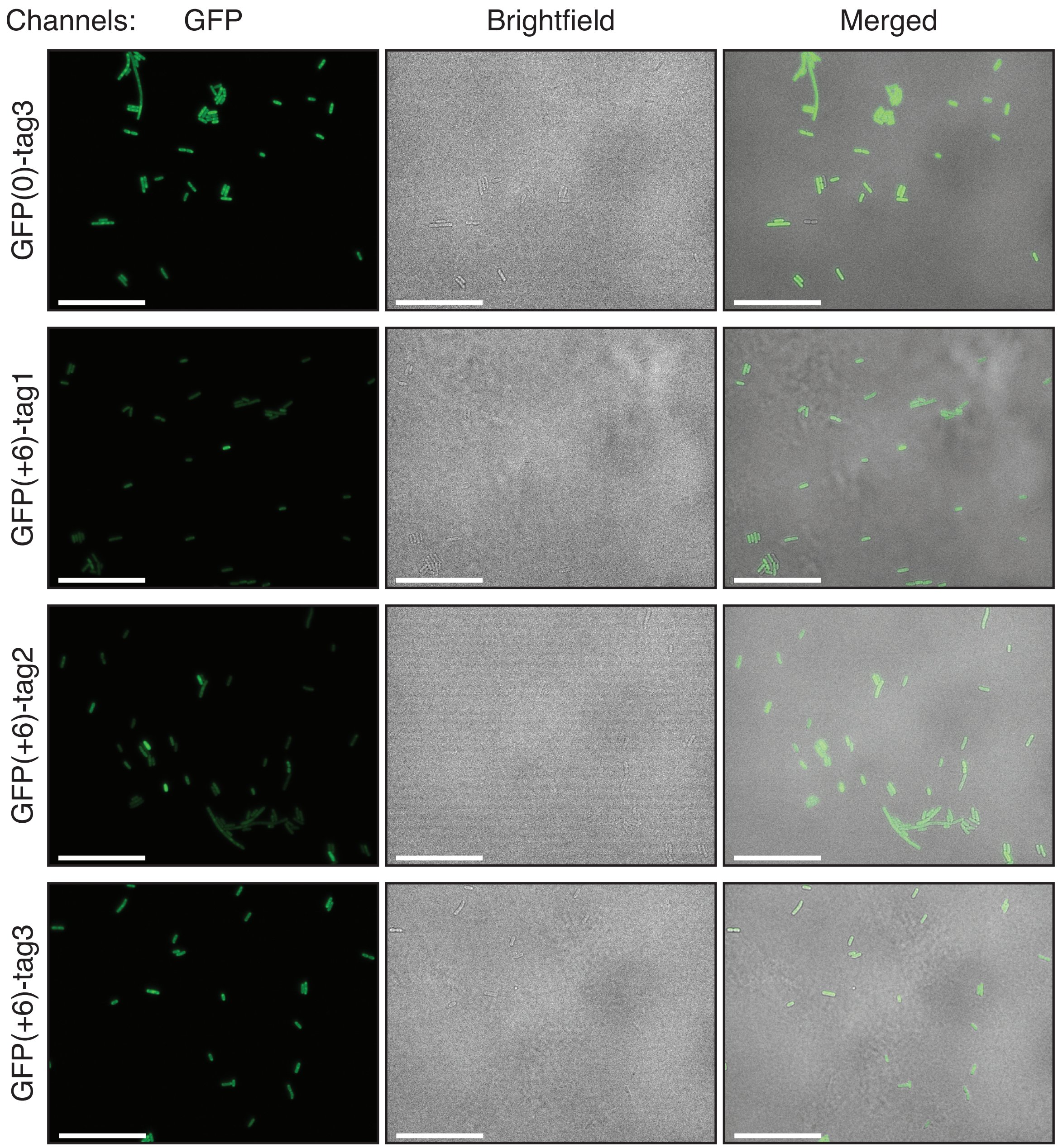

**Supplementary Figure 2. Representative microscopy images of *E. coli* cells expressing each GFP variant at 2 h post-induction.** GFP, brightfield, and merged channels depict a population of cells for each variant. Scale bars are 25 $\mu$m.

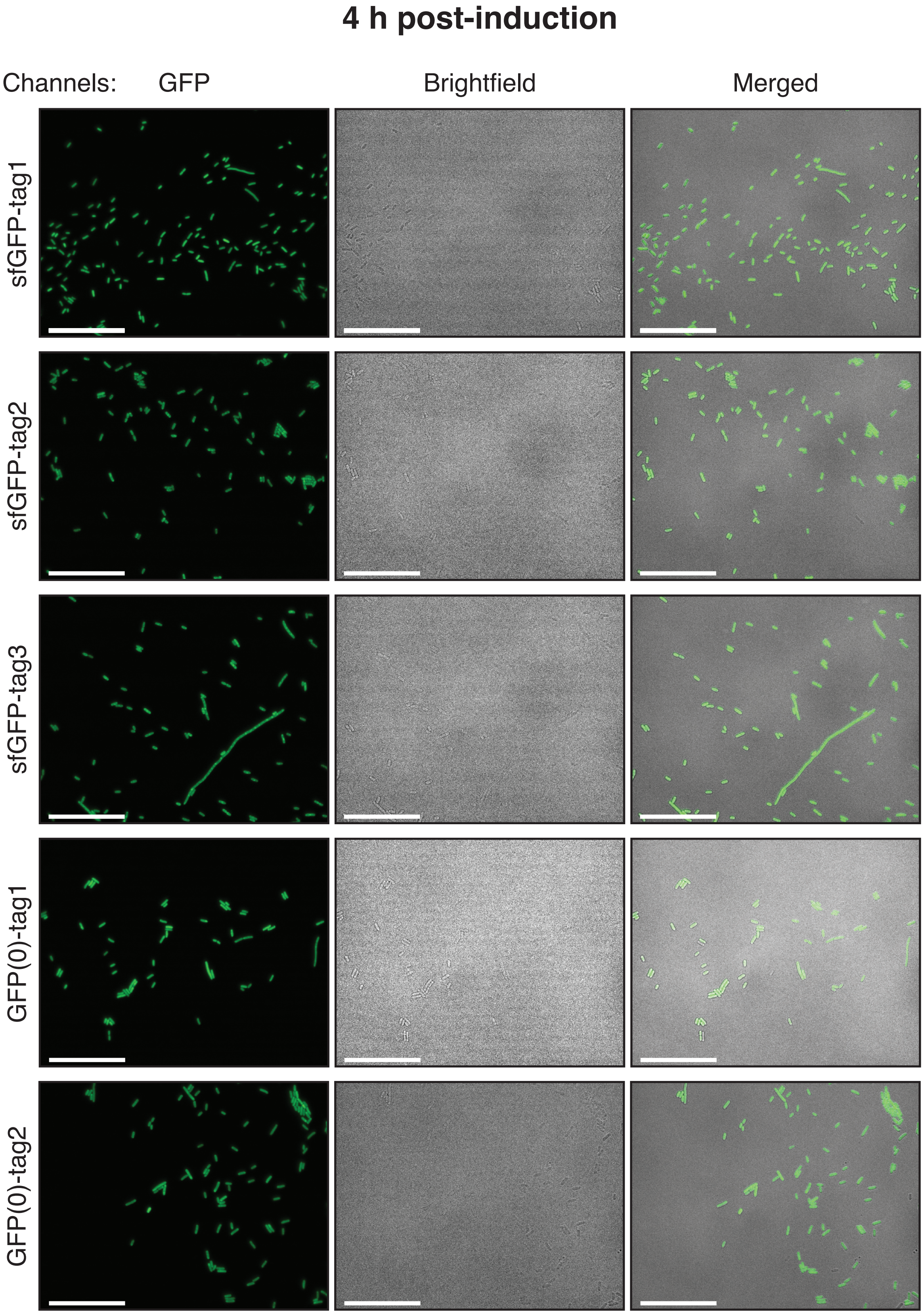
4 h post-induction
Channels:
GFP
Brightfield
Merged
sfGFP-tag1
sfGFP-tag2
sfGFP-tag3
GFP(0)-tag1
GFP(0)-tag2

### 4 h post-induction (continued)

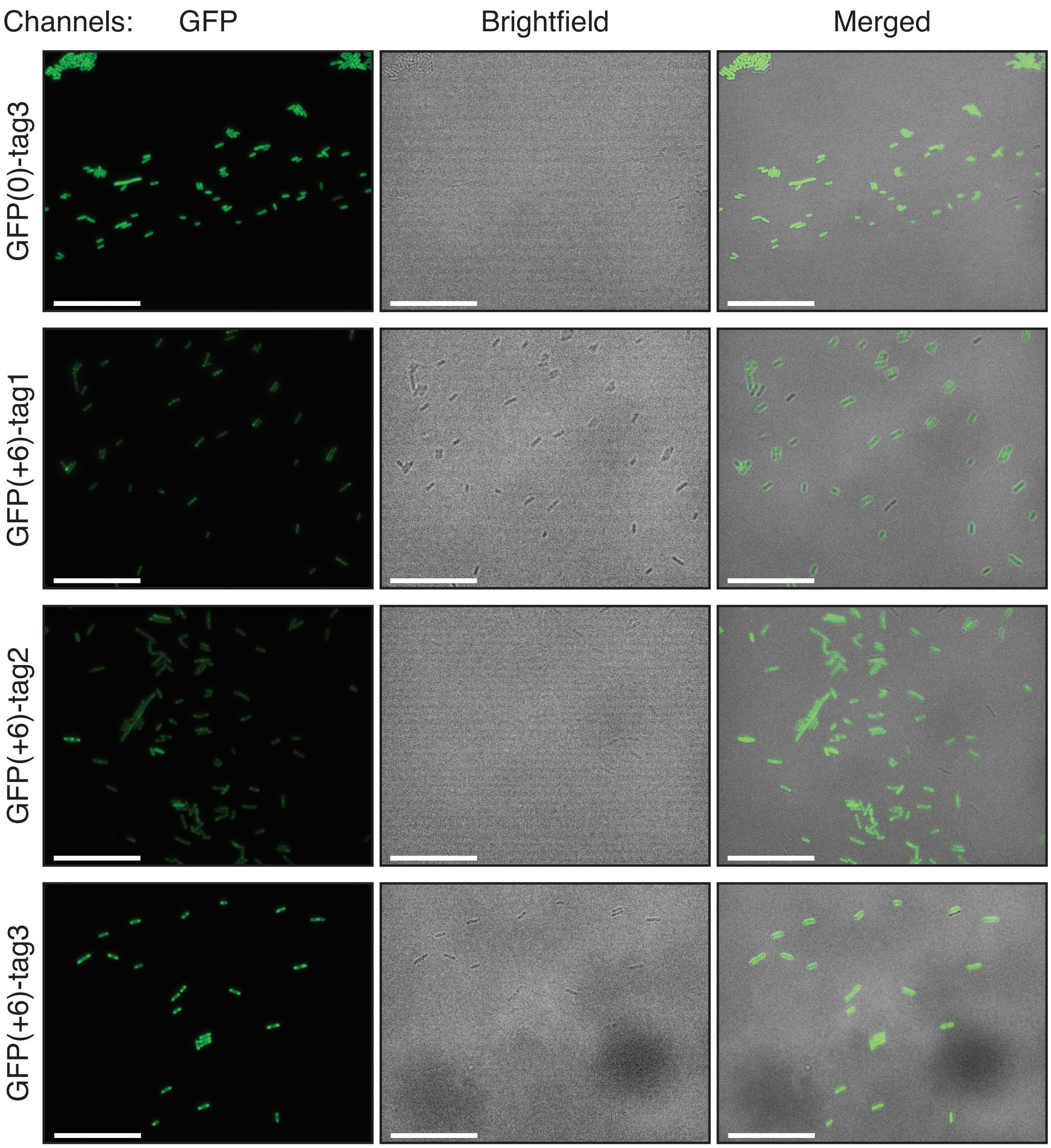

**Supplementary Figure 3. Representative microscopy images of *E. coli* cells expressing each GFP variant at 4 h post-induction.** GFP, brightfield, and merged channels depict a population of cells for each variant. Scale bars are 25 $\mu$m.

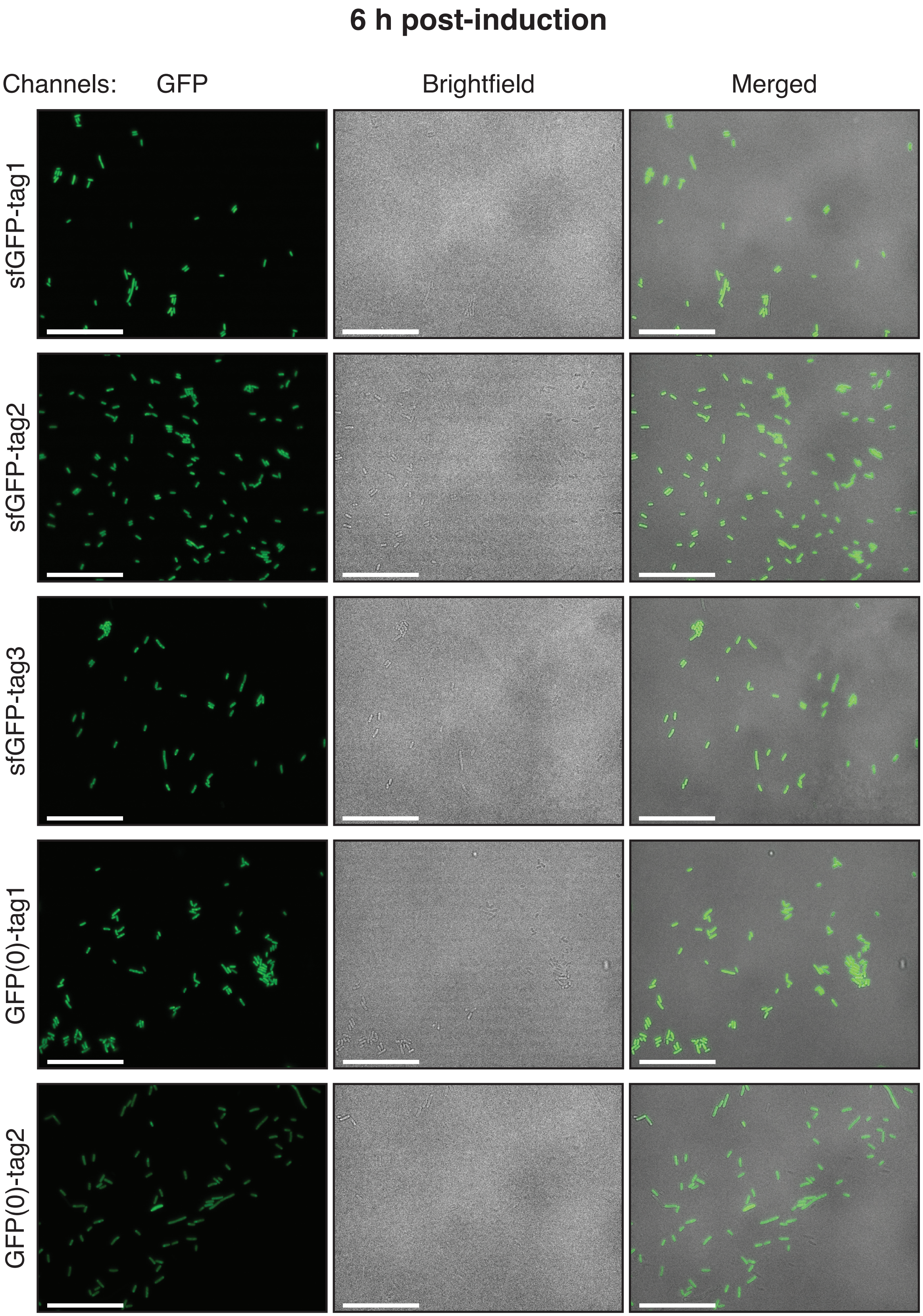
6 h post-induction
Channels:
GFP
Brightfield
Merged
sfGFP-tag1
sfGFP-tag2
sfGFP-tag3
GFP(0)-tag1
GFP(0)-tag2

### 6 h post-induction (continued)

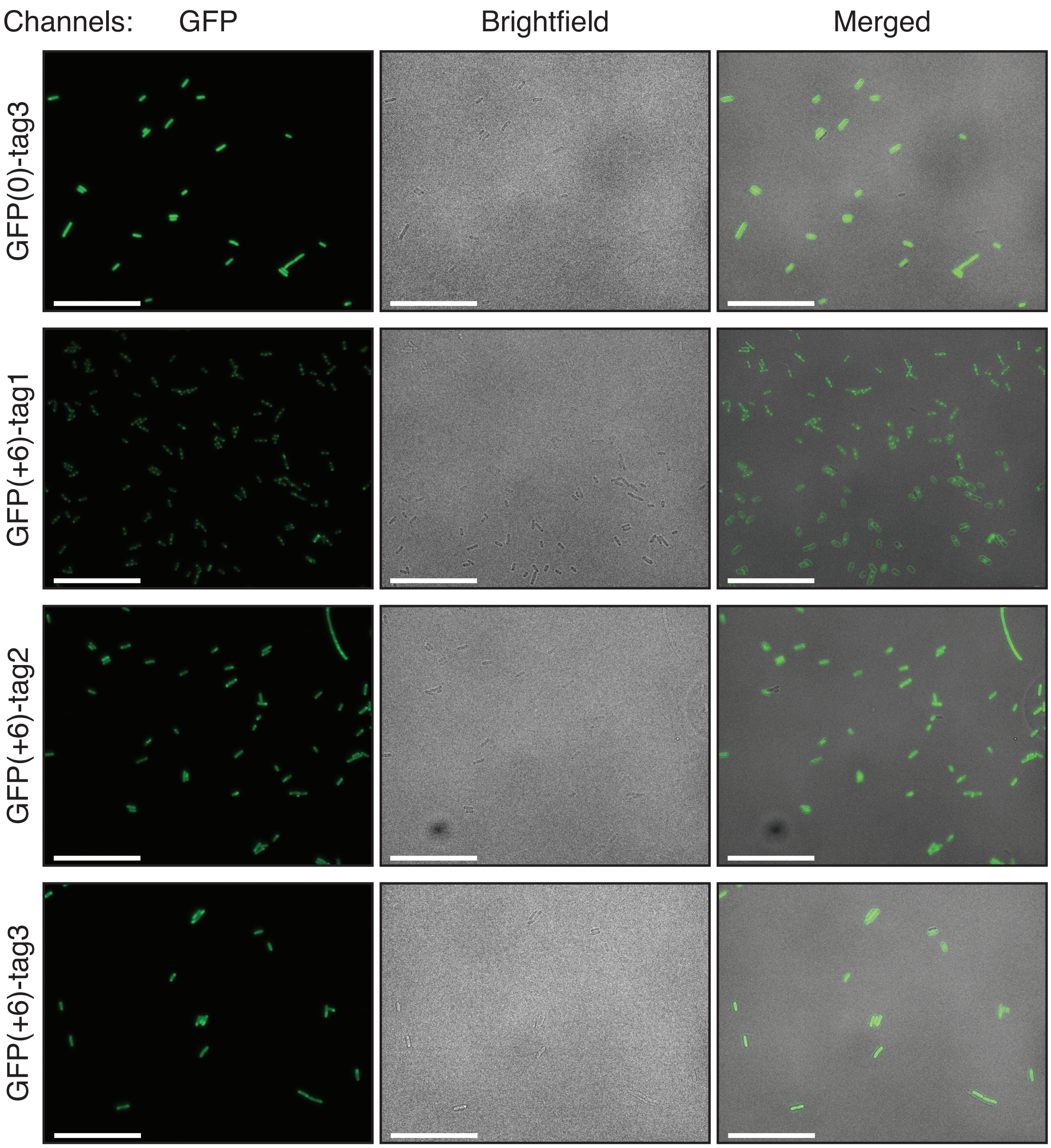

**Supplementary Figure 4. Representative microscopy images of *E. coli* cells expressing each GFP variant at 6 h post-induction.** GFP, brightfield, and merged channels depict a population of cells for each variant. Scale bars are 25 $\mu$m.

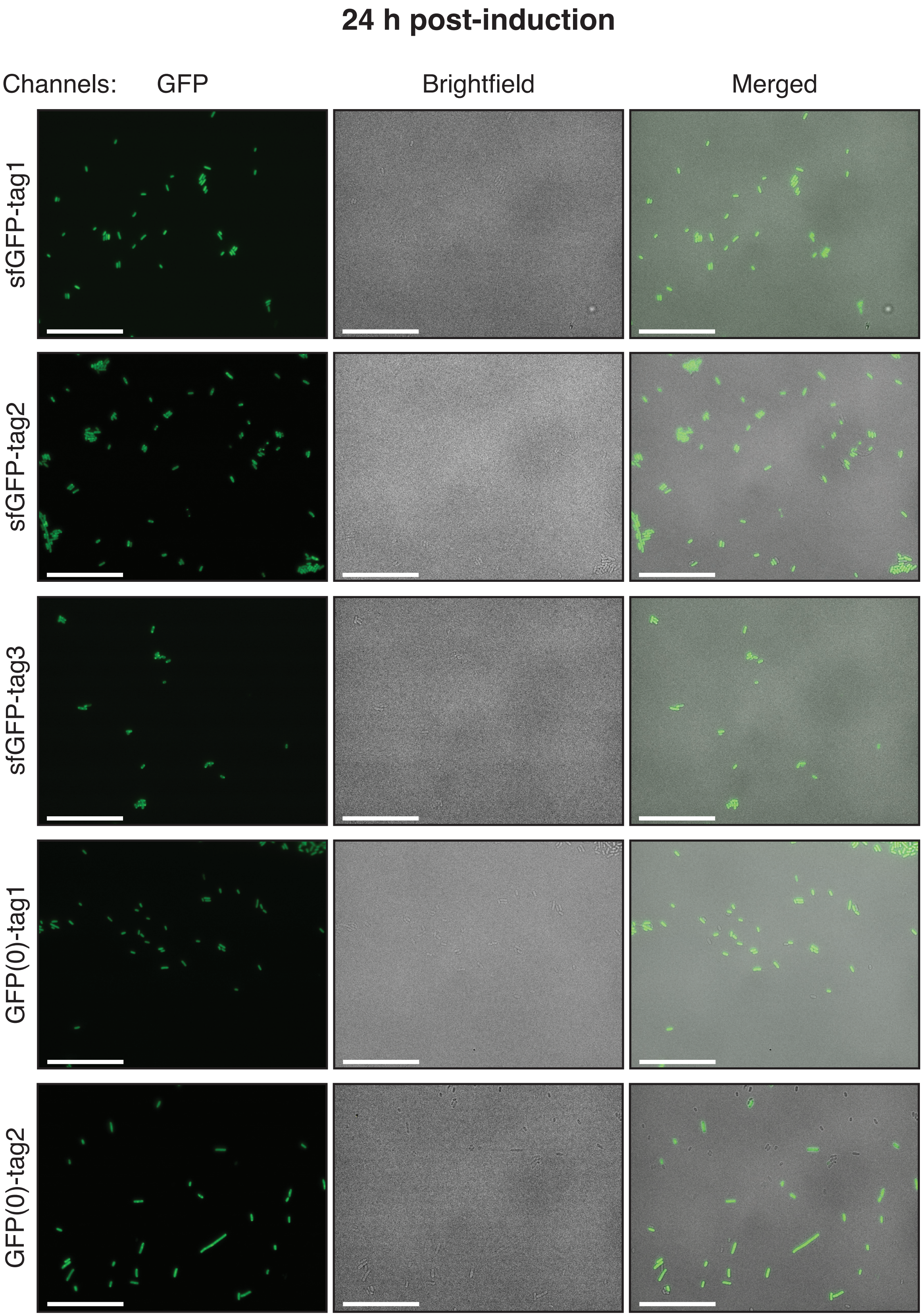
24 h post-induction
Channels:
GFP
Brightfield
Merged
sfGFP-tag1
sfGFP-tag2
sfGFP-tag3
GFP(0)-tag1
GFP(0)-tag2

### 24 h post-induction (continued)

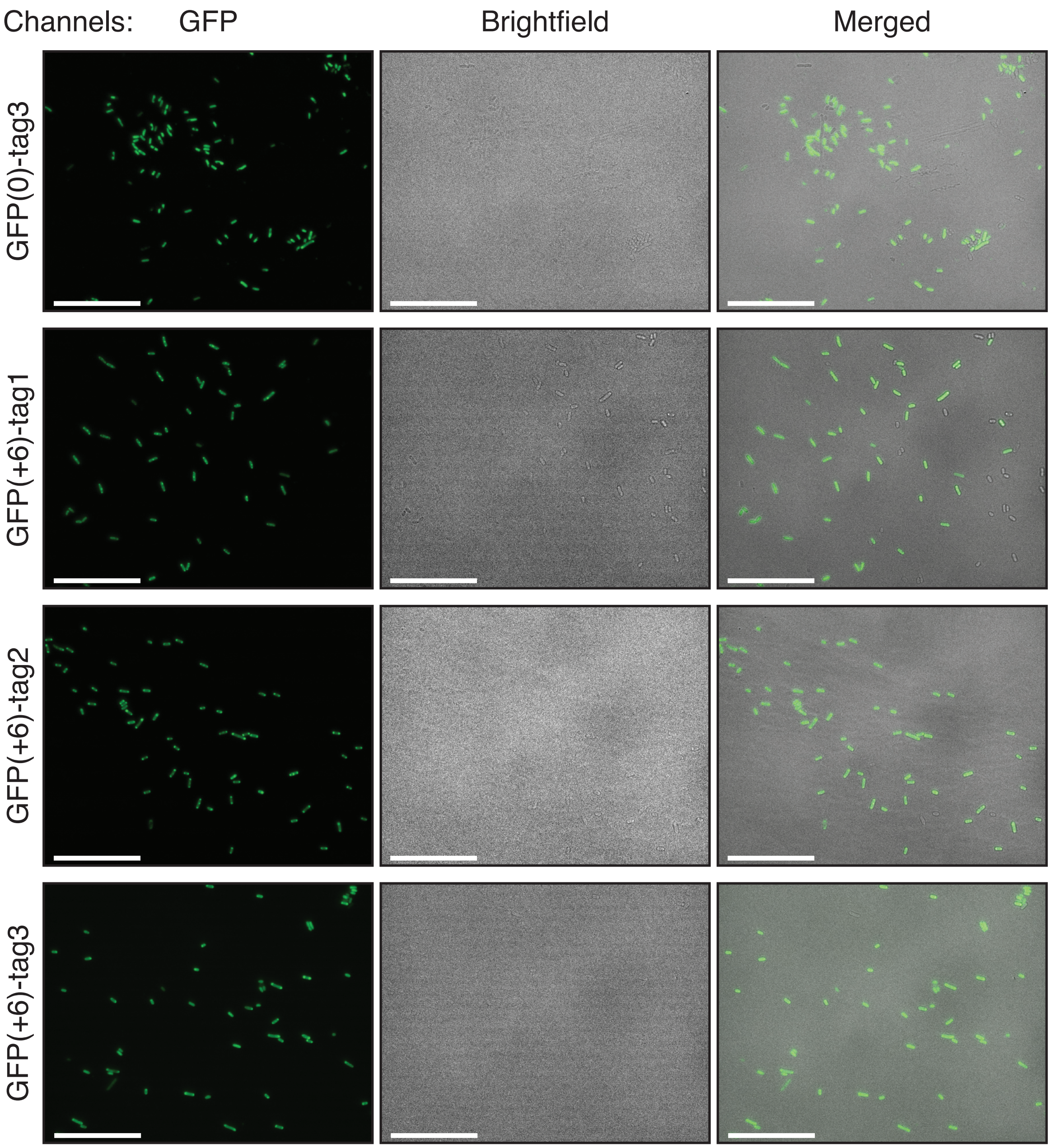

**Supplementary Figure 5. Representative microscopy images of *E. coli* cells expressing each GFP variant at 24 h post-induction.** GFP, brightfield, and merged channels depict a population of cells for each variant. Scale bars are 25 $\mu$m.

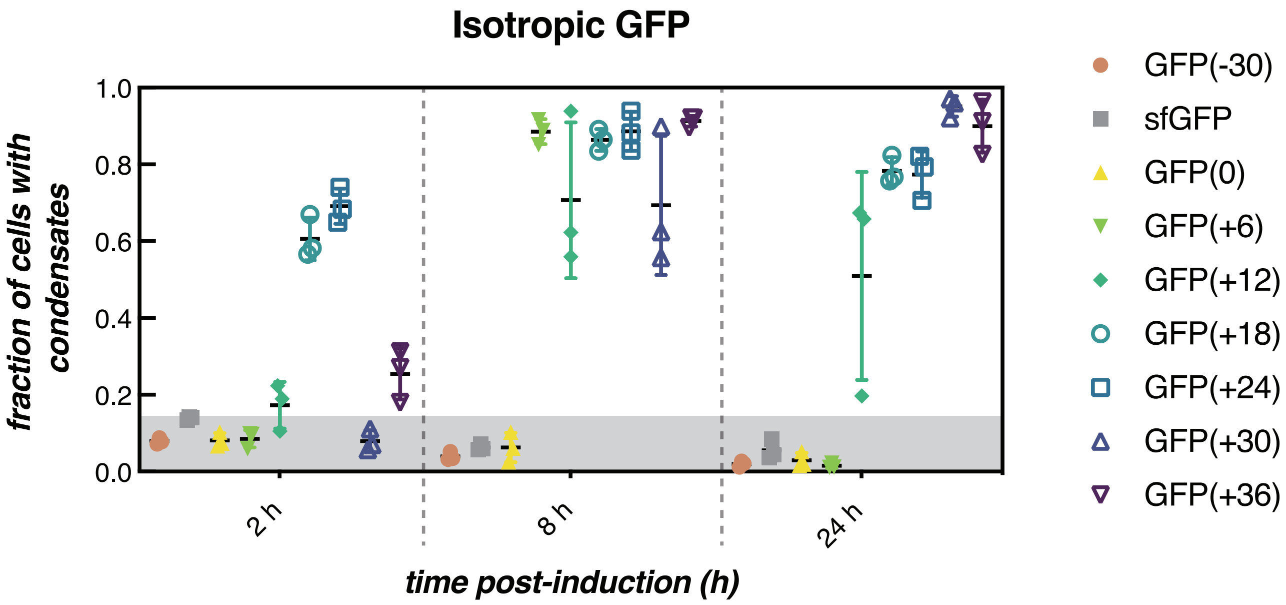

**Supplementary Figure 6. Supercationic isotropic GFP variants form condensates over time.** Image analysis further confirms intracellular phase separation in cells that express GFP variants with sufficient positive charge. As noted preivously, GFP variants with net charge ≥ +12 form and maintain condensates as protein is expressed post induction [5]. GFP(+6) undergoes reversible intracellular phase separation. The fracion of cells with condensates for three biological replicates (data points), the average (black line) and standard deviation (error bar) are plotted for each variant.

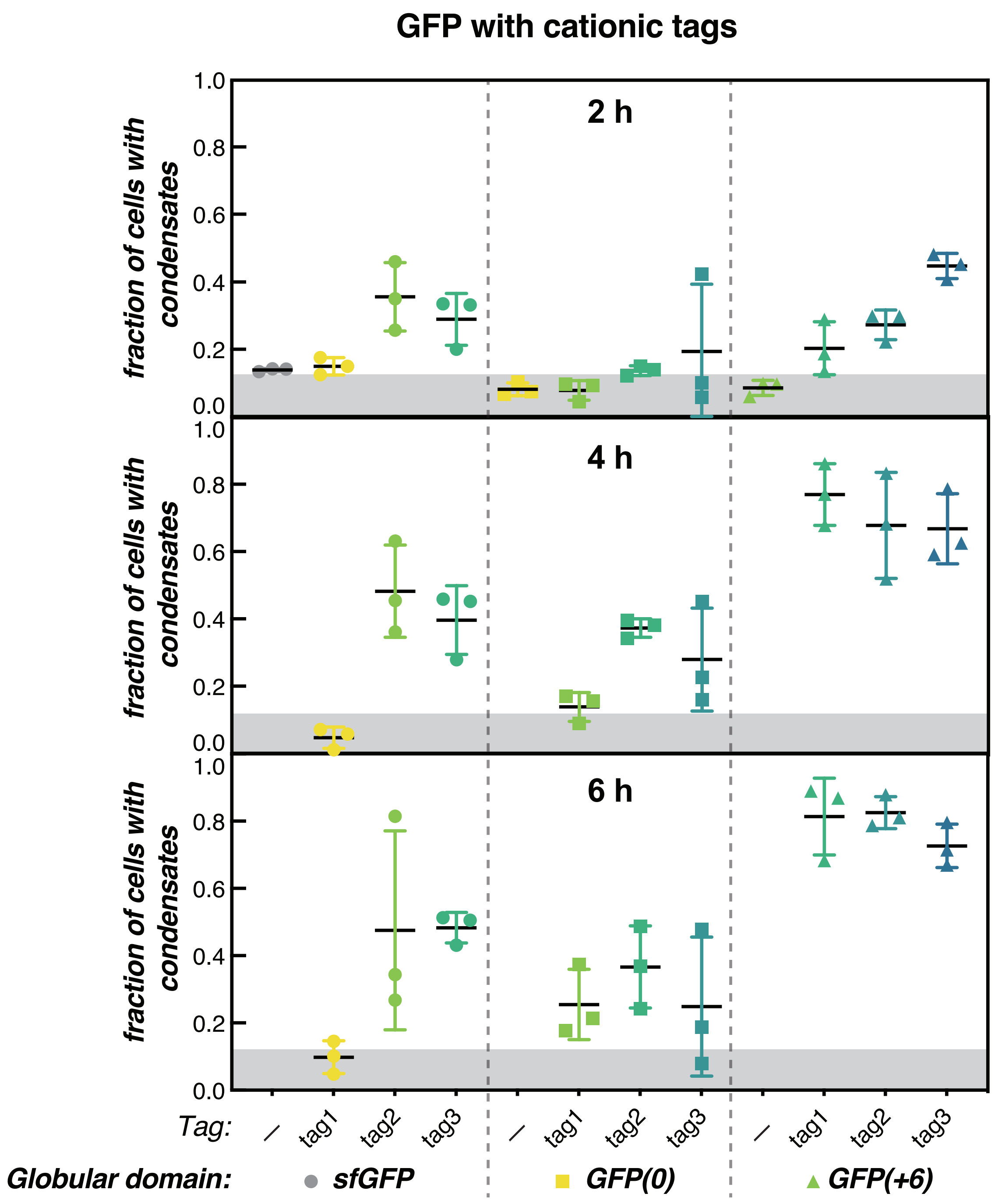

**Supplementary Figure 7. Tagged GFP variants can form condensates over time.** Quantification of microscopy images (n ≥ 416 total cells for each variant) at 2, 4, and 6 h post-induction. Data for isotropic variants were collected at 2 h, but not at 4 or 6 h [5]. The fraction of cells that contained detectable condensates were plotted for each variant. Generally, variants with a fraction < 0.11 did not form observable condensates. The average and standard deviation of three biological replicates are plotted for each variant.

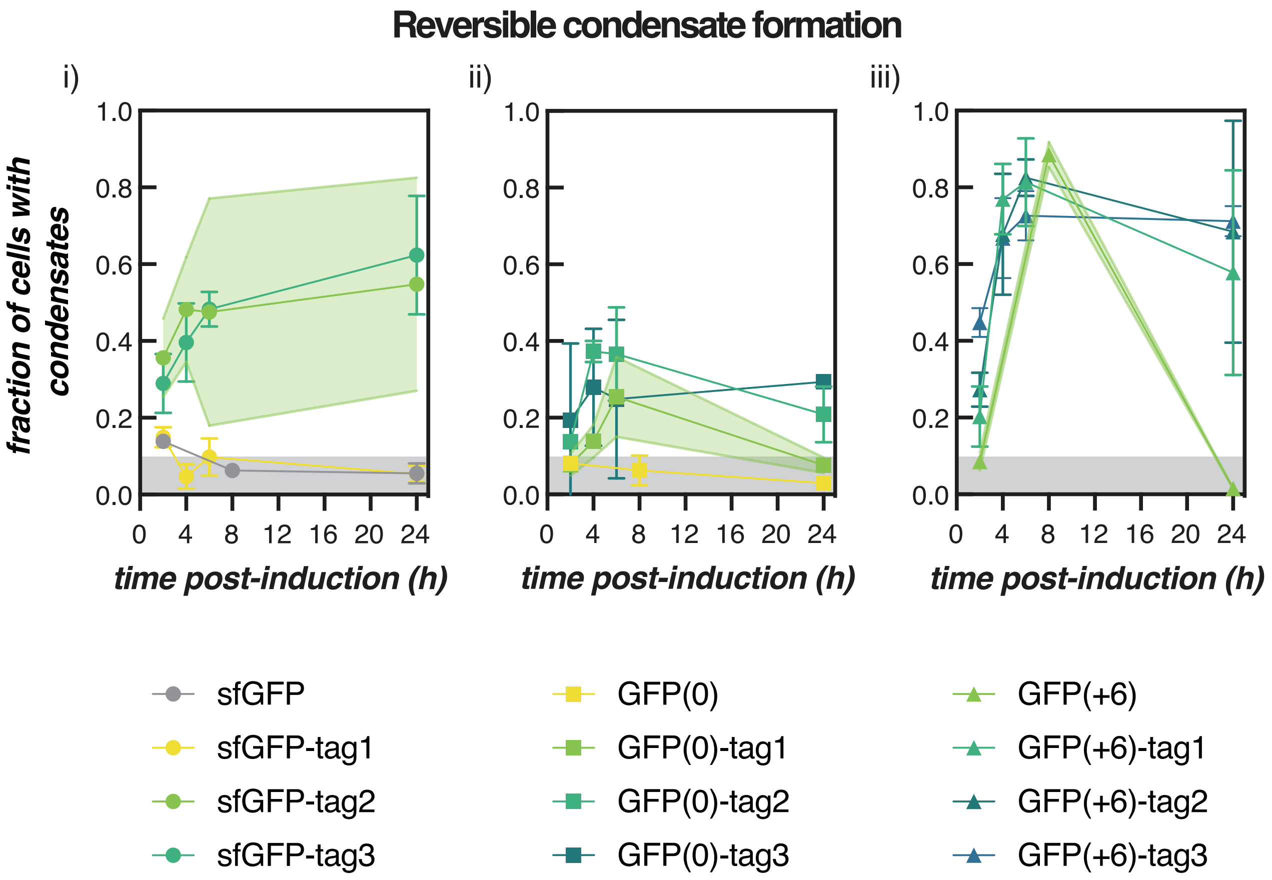

**Supplementary Figure 8. Reversible condensate formation.** The results from Supplementary Fig. 7 were replotted as a function of time instead of by protein variant. The gray shaded region represents an approximate threshold, below which cells did not form condensates. Condensate formation was observed for cells expressing GFP(0)-B1 (ii) and GFP(+6) (iii) at intermediate time points (6 h and 8 h, respectively). However, these condensates disappeared at 24 h, suggesting that condensate formation is reversible for these variants. The average and standard deviation of three biological replicates are plotted for each variant.

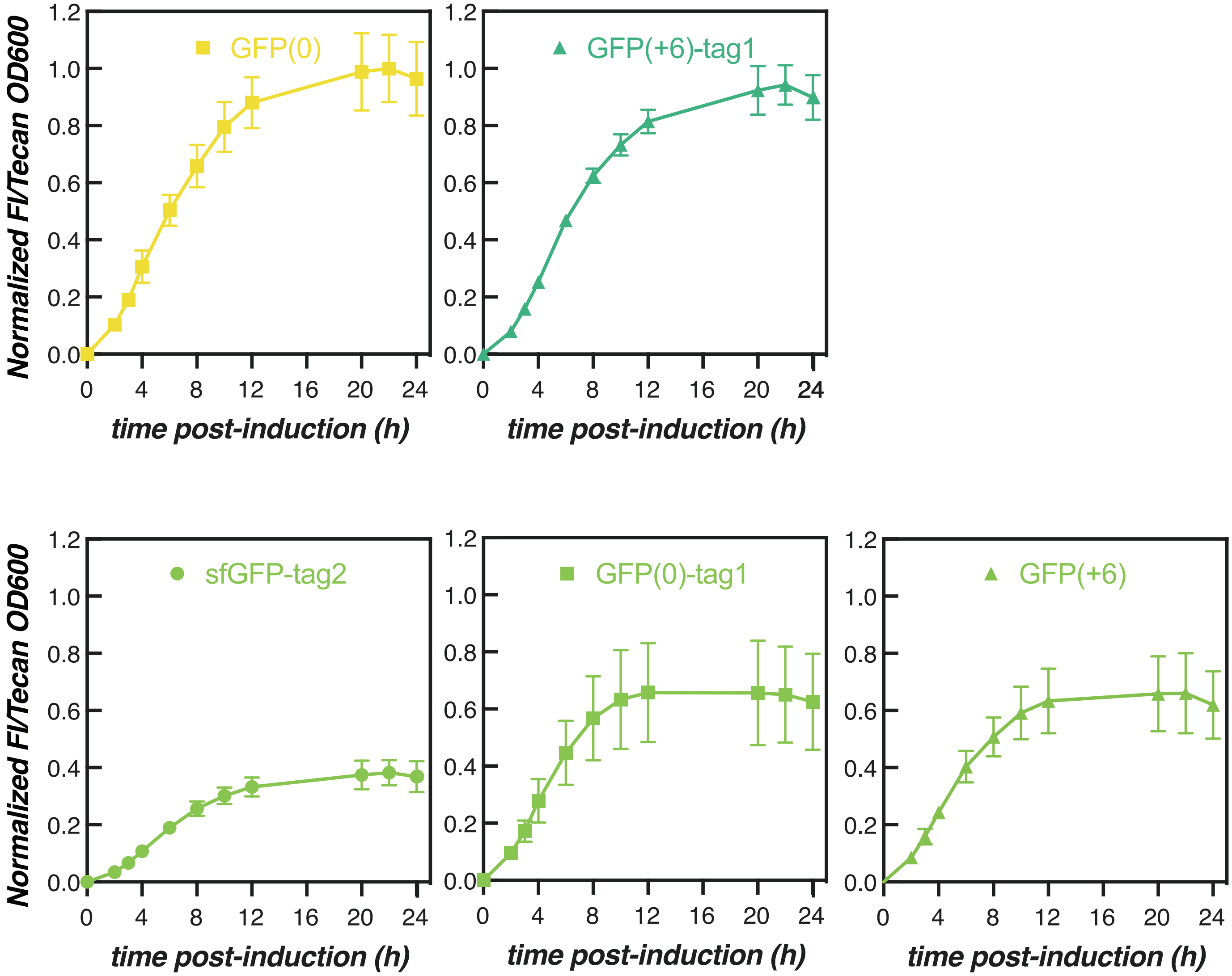

**Supplementary Figure 9. Cell growth and protein expression.** The ratio of GFP fluorescence intensity to cell density was plotted on separate graphs for select GFP variants. This ratio was used as a proxy for the intracellular GFP concentration. The intracellular concentrations of like-charged GFP(+6) and GFP(0)-tag1 were similar, providing one explanation for how these two strains were able to traverse the phase boundary. Plotted are the average and standard deviation of the normalized fluorescence intensity/Tecan OD ratio for each variant. This ratio was calculated by averaging across three biological replicates and then normalizing to the highest average fluorescence intensity/Tecan OD ratio amongst all GFP variants (GFP(0)).

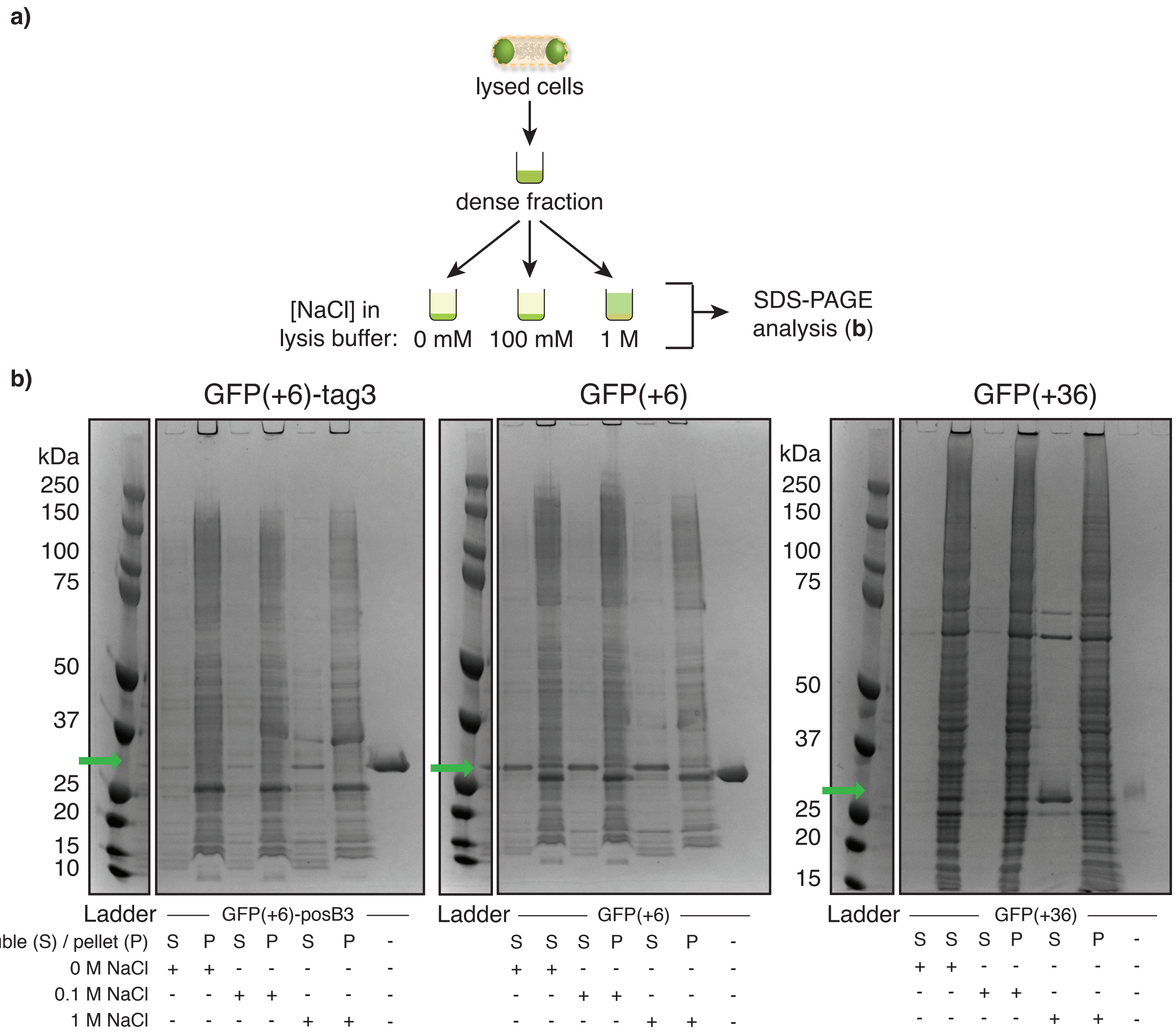

**Supplementary Figure 10. Solubility of GFP(+6)-tag3, GFP(+6), and GFP(+36) in buffers of varied ionic strength.** (a) Schematic depicting the extraction of GFP from the cell as previously reported [5]. Briefly, lysed cells were separated into soluble and insoluble fractions by centrifugation. The insoluble fraction was then washed and resuspended in a buffer containing different concentrations of NaCl. These samples were separated into their soluble and insoluble fractions, which were collected and analyzed on SDS-PAGE gels. (b) SDS-PAGE gels depicting similarities in solubility between the GFP(+6)-tagged variant and a coacervate-forming isotropic GFP variant, GFP(+36). "S" refers to the soluble fraction, and "P" refers to the pellet. "+" indicates treatment with buffers of varying NaCl concentration (0 M, 0.1 M, or 1 M), Precision Plus Protein Dual Color standard (Bio-rad) was loaded in lane 1 for all gels. Lanes 2-7 depict different buffer treatments of cell lysates and lane 8 corresponds to the respective purified GFP. The ladders shown were loaded onto the same gels as the individual GFP variants; however, irrelevant lanes were excluded for clarity. The band corresponding to GFP is indicated by ⟶ .

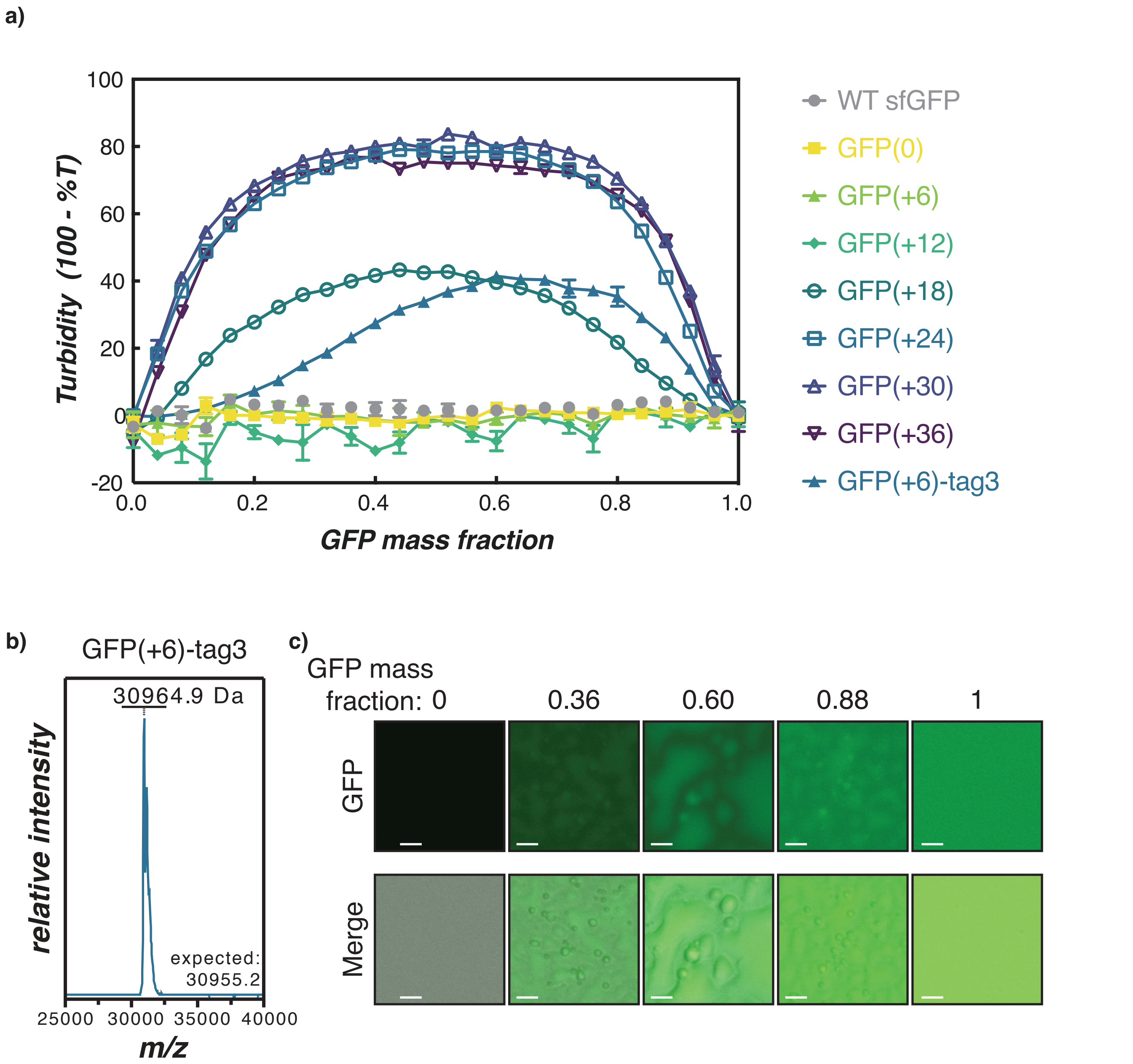

**Supplementary Figure 11.** ***In vitro*** **complex coacervation of RNA with GFP(+6)-tag3.** (a) Turbidity assay of supercharged cationic GFP variants with total RNA from torula yeast as reported previously [5]. The turbidity of the tagged variant falls between the untagged control, GFP(+6), and the isotropic protein with equivalent charge, GFP(+24). The average and standard deviation of three technical replicates are plotted for each variant. (b) MALDI-TOF spectrum of purified GFP(+6)-tag3. The expected molecular weight was calculated based on the primary amino acid sequence without the methionine from the start codon. (c) Microscopy images of GFP(+6)-tag3/RNA mixtures at 1 mg/mL total macromolecule concentration. Mixtures were prepared in a physiological buffer that mimics the intracellular ion concentration in *E. coli* (70 mM $K_2HPO_4$, 60 mM KCl, 40 mM NaCl, pH 7.4). Scale bar is 10 $\mu$M.

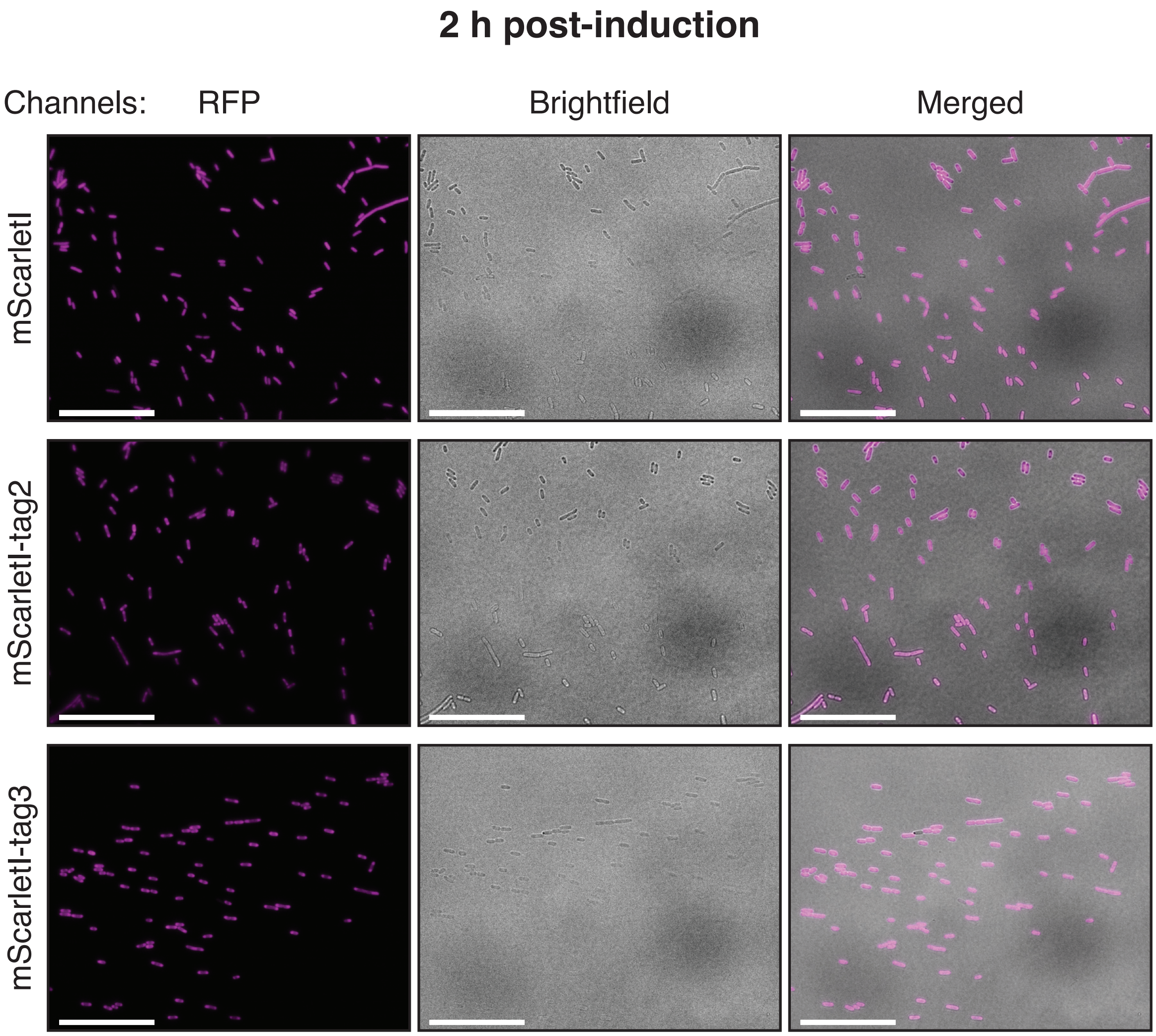

**Supplementary Figure 12. Representative microscopy images of *E. coli* cells expressing each mScarletI variant at 2 h post-induction.** TexasRed, brightfield, and merged channels depict a population of cells for each variant. Scale bars are 25 $\mu$m.

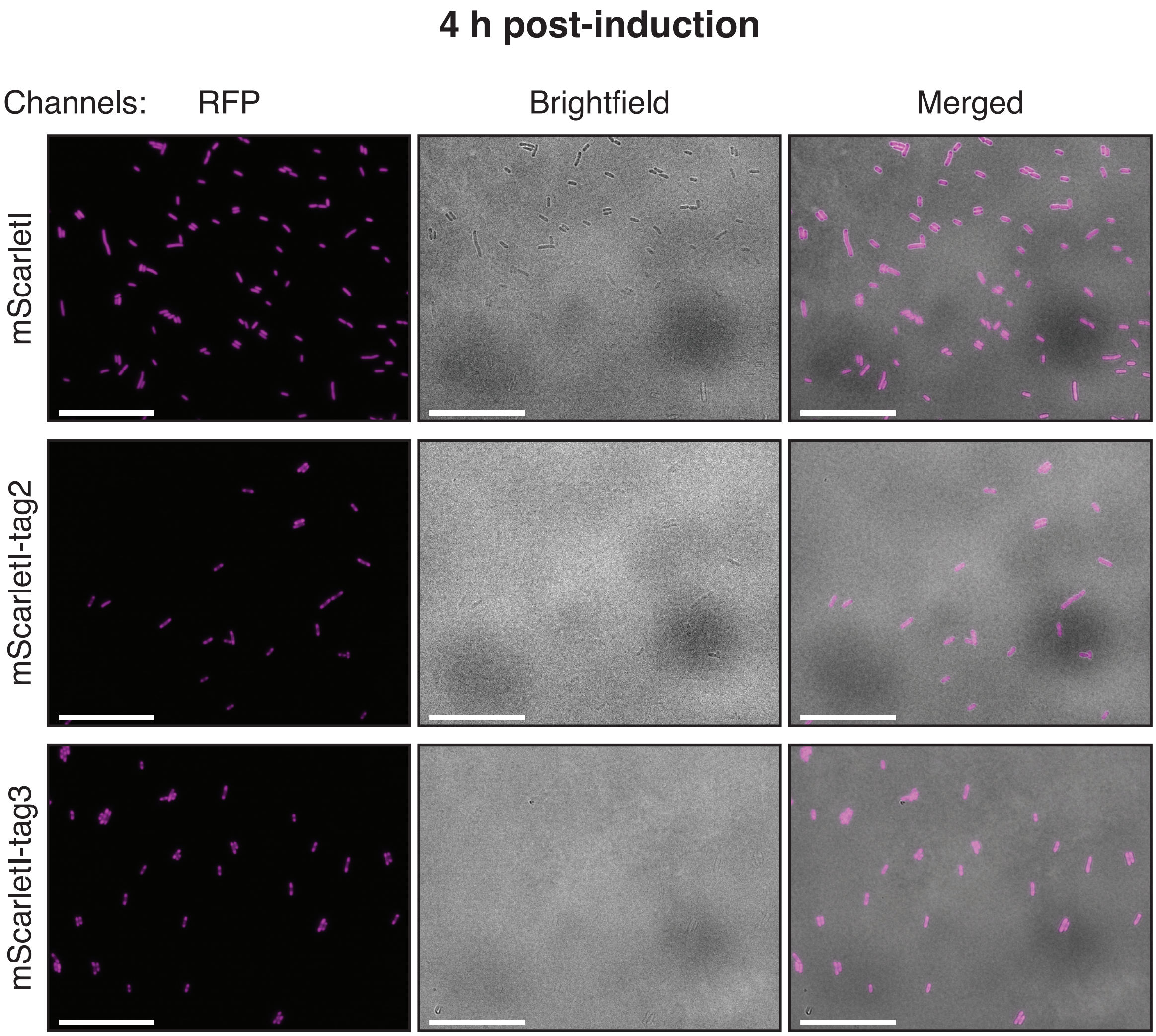

**Supplementary Figure 13. Representative microscopy images of *E. coli* cells expressing each mScarletI variant at 4 h post-induction.** TexasRed, brightfield, and merged channels depict a population of cells for each variant. Scale bars are 25 $\mu$m.

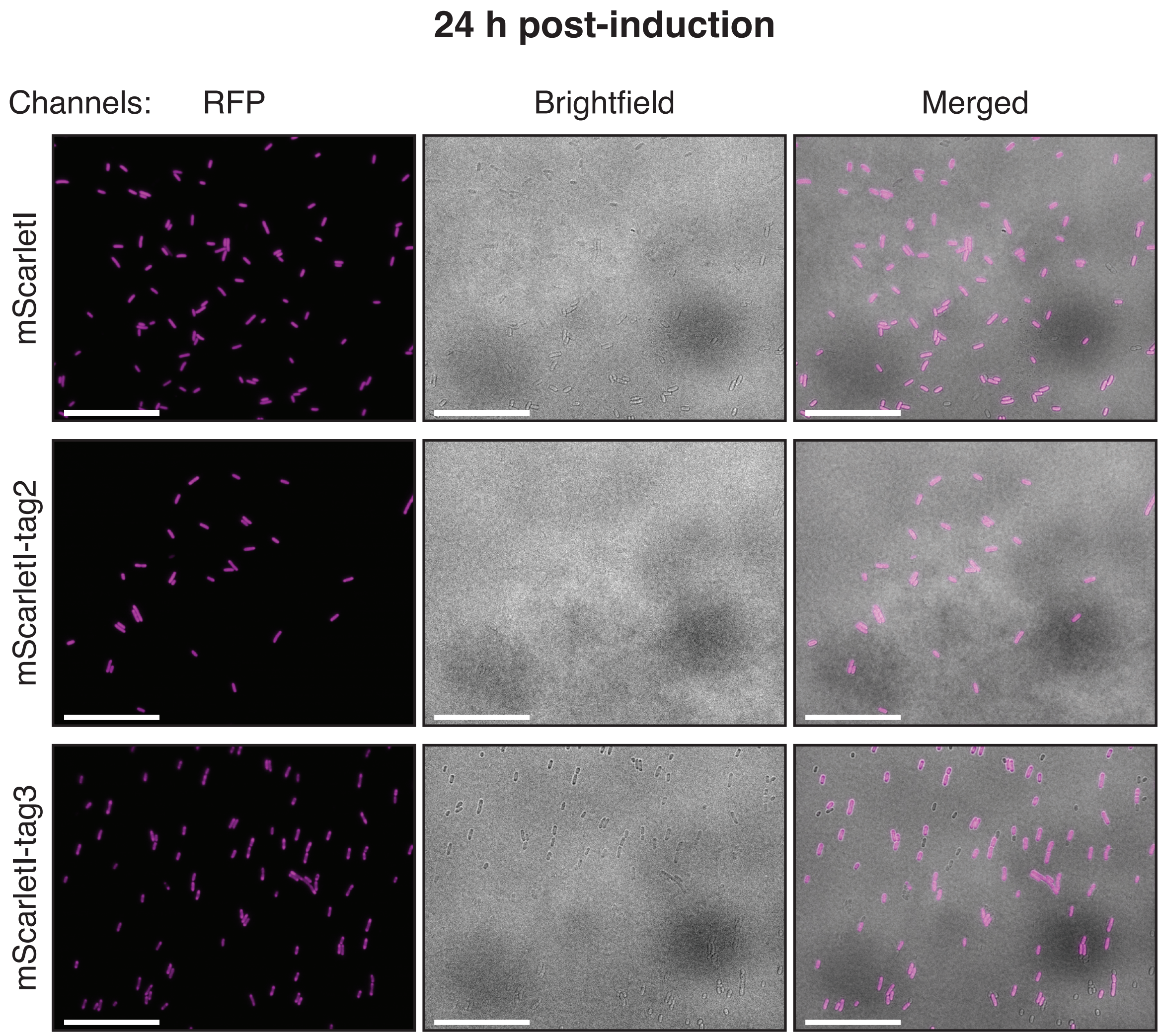

**Supplementary Figure 14. Representative microscopy images of *E. coli* cells expressing each mScarletI variant at 24 h post-induction.** TexasRed, brightfield, and merged channels depict a population of cells for each variant. Scale bars are 25 $\mu$m.

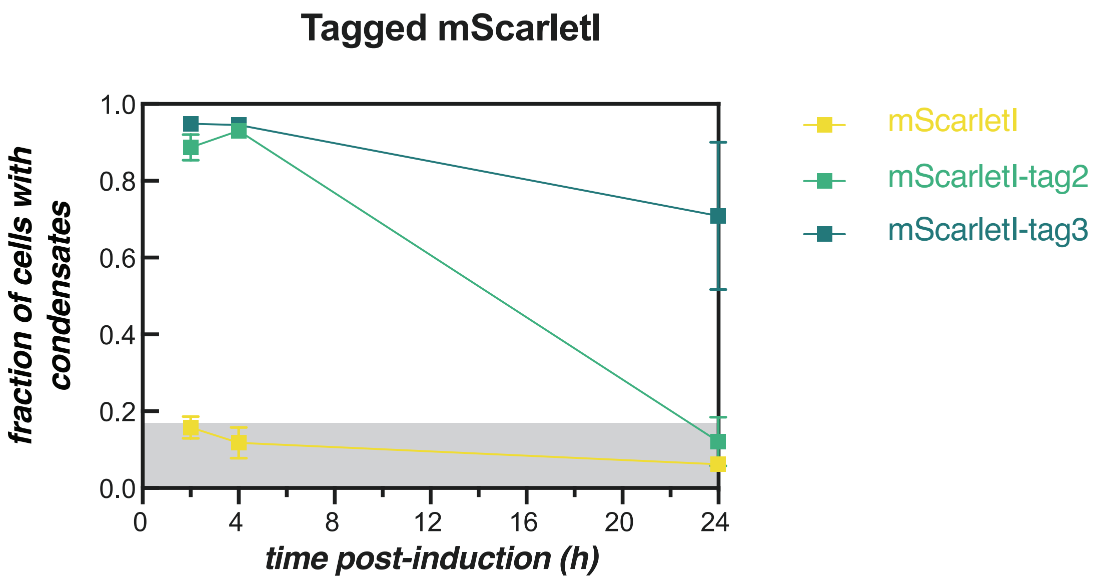

**Supplementary Figure 15. Supercationic mScarletI variants form condensates over time.** Quantificaition of microscopy images taken at 2, 4, and 24 h post-induction. The fraction of cells that contained detectable condensates were plotted for each variant as a function of time post-induction. This analysis further confirmed intracellular phase separation in cells that expressed mScarletI variants with sufficient positive charge. The average and standard deviation of three biological replicates are plotted for each variant.

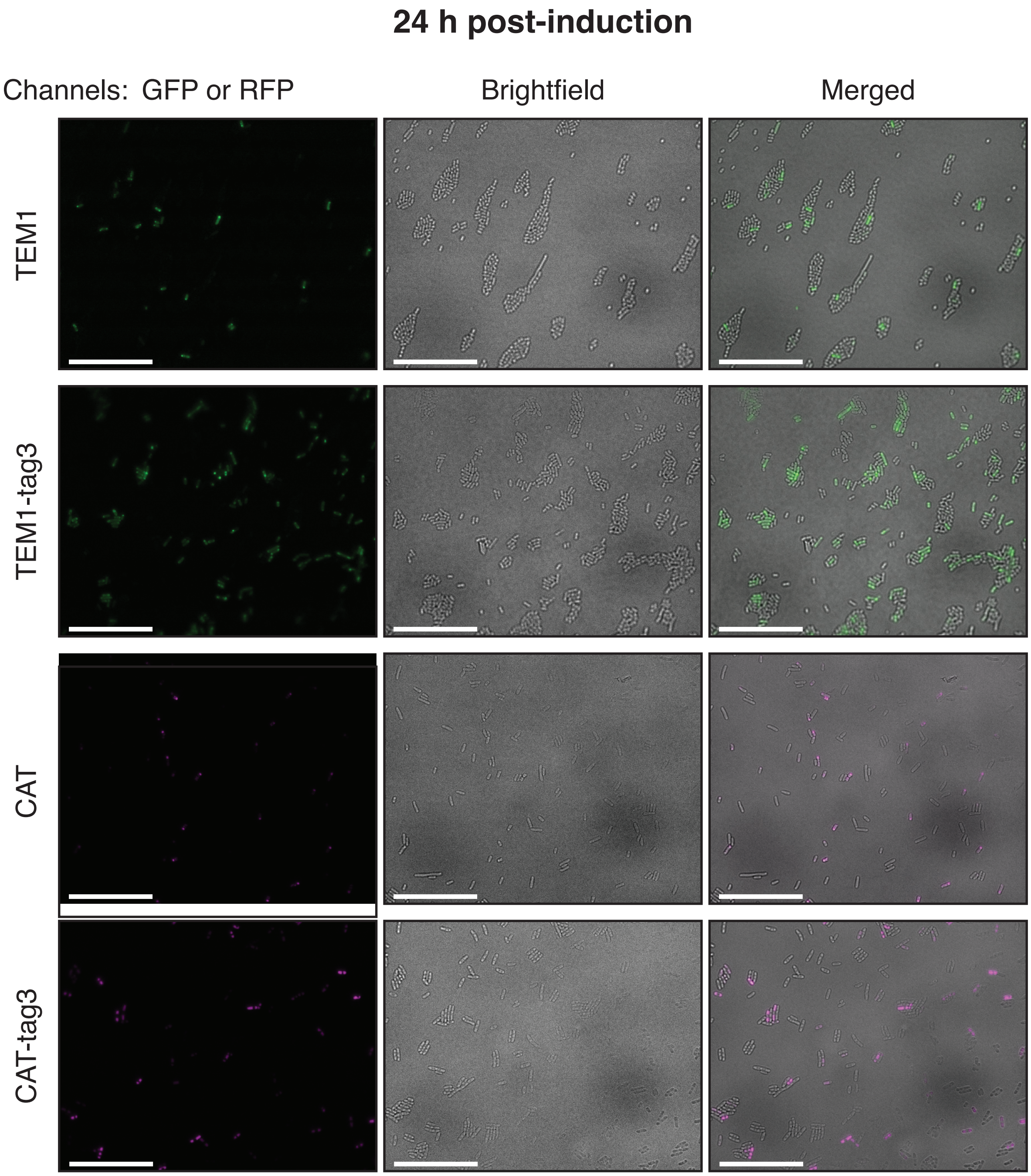

**Supplementary Figure 16. Representative microscopy images of *E. coli* cells expressing beta-lactamase (TEM1) or chloramphenicol acetyltransferase (CAT) variants at 24 h post-induction.** Beta-lactamase variants were stained with 10 $\mu$M FlAsH-$EDT_2$ and images of cell populations were acquired on GFP and brightfield channels. CAT variants were stained with 1 $\mu$M ReAsH-$EDT_2$ and images of cell populations were acquired on TexasRed and brightfield channels. Scale bars are 25 $\mu$m.

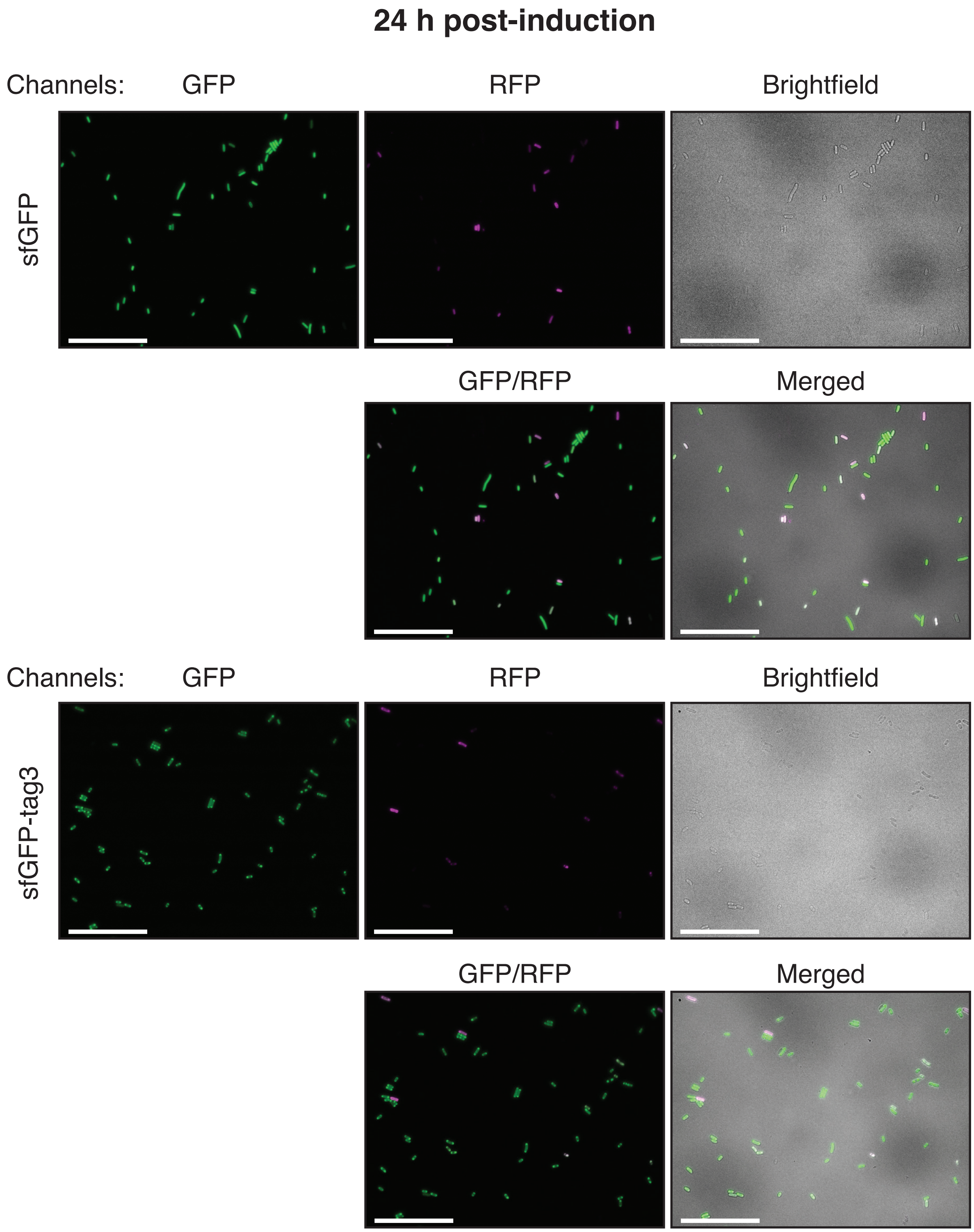

**Supplementary Figure 17. Representative microscopy images of *E. coli* cells expressing sfGFP or sfGFP-tag3 containing an N-terminal TC tag at 24 h post-induction.** Cells expressing sfGFP and sfGFP-tag3 were stained with 1 $\mu$M ReAsH-EDT$_2$ and images of cell populations were acquired on GFP, TexasRed, and brightfield channels. Scale bars are 25 $\mu$m.

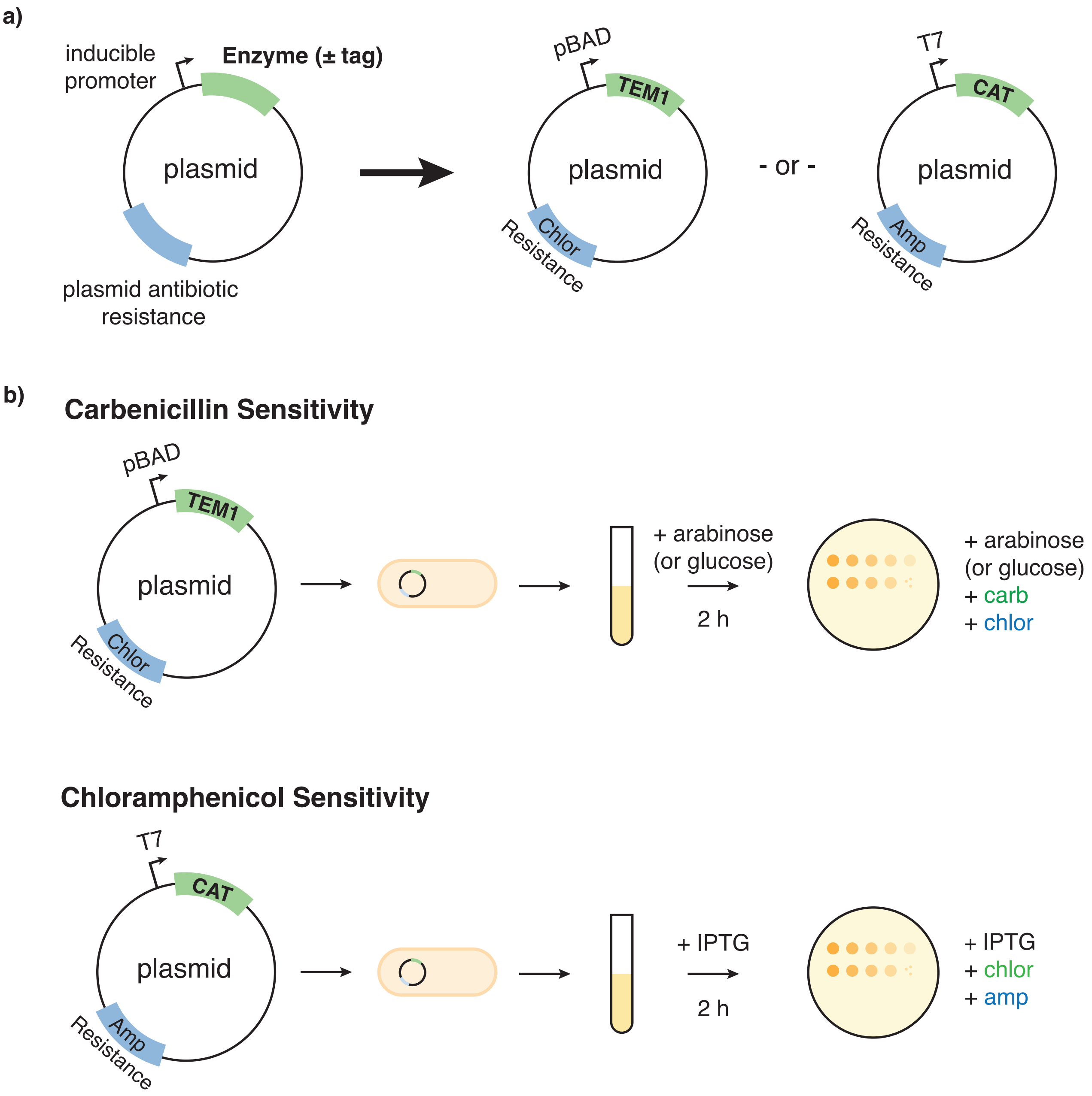

**Supplementary Figure 18. Experimental design of carbenicillin and chloramphenicol sensitivity assays.** (a) Design of plasmids expressing the beta-lactamase (TEM1) and chloramphenicol acetyltransferase (CAT) variants (in green). TEM1 and CAT expression were induced either by arabinose (pBAD) or IPTG (T7). Plasmids also contained a separate antibiotic resistance marker (in blue) to ensure plasmid retention during the sensitivity assays. (b) Spotting assays were conducted for beta-lactamase (TEM1) and chloramphenicol acetyltransferase (CAT) variants. Briefly, cells from overnight cultures were grown in fresh LB media and induced for 2 h to allow for condensate formation prior to spotting on LB agar plates containing the respective inducers and antibiotics. Plates were supplemented with varying concentrations of carbenicillin or chloramphenicol (in green) to test the sensitivity of the TEM1 or CAT variants, respectively. Plates were also supplemented with a 25 $\mu$g/mL chloramphenicol or 100 $\mu$g/mL ampicillin for plasmid selection (in blue).

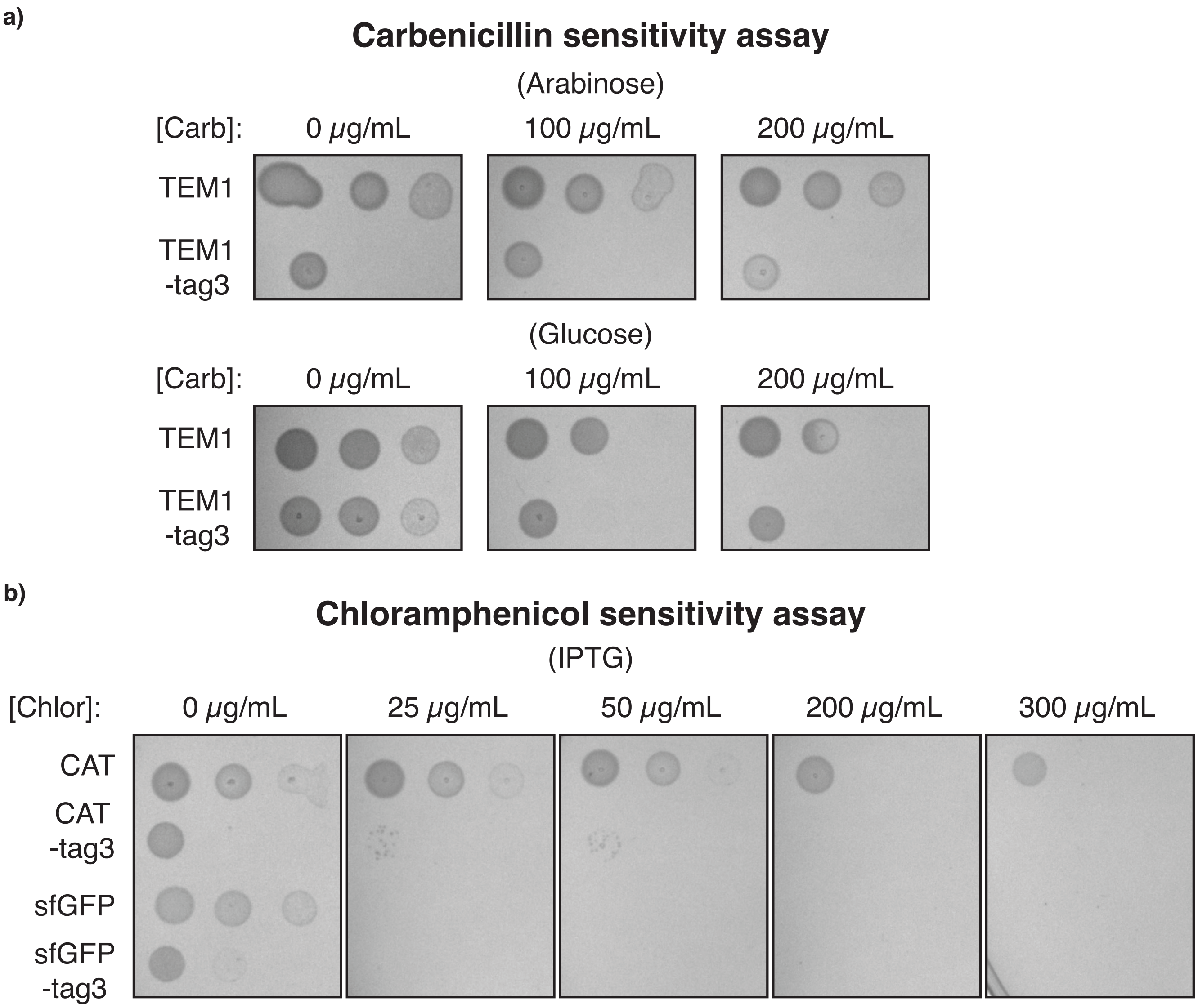

**Supplementary Figure 19. Carbenicillin and chloramphenicol sensitivity assays.** (a) TEM1 variants were grown on plates supplemented with either arabinose or glucose along with carbenicillin. Starting $OD_{600}$ ~ 1 in the leftmost column, followed by subsequent dilutions (20X and 400X are shown). Reduced growth on the control (no carbenicillin) arabinose plate suggested that condensate formation in the TEM1-tag3 variant reduced cell fitness but did not abolish beta-lactamase activity as these variants could grow on plates supplemented with increasing concentrations of chloramphenicol. Growth of TEM1-tag3 variants on glucose plates showed that TEM1-tag3 maintained enzymatic activity when exposed to carbenicillin despite suppressed expression. Minimal growth differences between TEM1 and TEM1-tag3 were observed in the absence of carbenicillin on glucose plates. (b) CAT variants were grown on plates supplemented with ITPG and chloramphenicol. sfGFP variants were also included as charge-equivalent, negative controls for antibiotic resistance. Starting $OD_{600}$ ~ 0.7 in the leftmost column, followed by subsequent dilutions (100X and 2000X are shown). Similar to the TEM1 variants, reduced fitness without elimination of enzyme activity was observed for tagged CAT variants.
